# An unprecedented small RNA-riboswitch interaction controls expression of a bifunctional pump that is essential for *Staphylococcus aureus* infection

**DOI:** 10.1101/2024.09.30.615890

**Authors:** Gabriela González-Espinoza, Karine Prévost, Fayyaz Hussain, Jana N. Radin, Carlos D. Vega Valle, Julie Maucotel, Marina Valente Barroso, Benoît Marteyn, Eric Massé, Pascale Romby, Thomas E. Kehl-Fie, Jens Georg, David Lalaouna

**Affiliations:** Université de Strasbourg, CNRS, Architecture et Réactivité de l’ARN, UPR9002, F-67000 Strasbourg, France; CRCHUS, RNA Group, Department of Biochemistry and Functional Genomics, Faculty of Medicine and Health Sciences, Université de Sherbrooke, 3201 Jean Mignault Street, Sherbrooke, QC J1E 4K8, Canada; University of Freiburg, Faculty of Biology, Institute of Biology III, Genetics and Experimental Bioinformatics, D-79104 Freiburg, Germany; Department of Microbiology and Immunology, Carver College of Medicine, University of Iowa, Iowa City, IA, USA

**Keywords:** *Staphylococcus aureus*, metal homeostasis, manganese, iron, small RNA, riboswitch, efflux pump, virulence, therapeutic target

## Abstract

Maintaining manganese and iron homeostasis is critical for the human pathogen *Staphylococcus aureus*. To counteract metal-based host defense strategies (e.g., nutritional immunity, metal poisoning), *S. aureus* uses a combination of metal-sensing transcription factors and regulatory RNAs to maintain metal homeostasis. In this study, we uncovered an unprecedented interaction between a cis- and a trans-acting regulatory RNA controlling a conditionally essential gene, *mntY*, encoding a Mn efflux pump. This broadly conserved RNA-RNA interaction between a Fe-responsive sRNA and a Mn-sensing riboswitch allows the integration of Fe- and Mn-related stresses, notably encountered at the infection site, to fine-tune *mntY* expression. Remarkably, deletion of the *mntY* gene is strongly pleomorphic, causing growth defects, altering virulence factor expression, immune evasion, and survival during infection. We demonstrated that MntY is critical for the adaptation of *S. aureus* to both low and high Mn environments, due to its dual role in metalation of Mn-dependent exoenzymes and Mn detoxification. These findings point to MntY as a promising new therapeutic target to combat multidrug-resistant staphylococcal infections.

## INTRODUCTION

Metals such as iron (Fe) and manganese (Mn) are essential nutrients for life due to their crucial roles in numerous structural and catalytic cellular processes^1,2^. During infection, the host takes advantage of this dependency by removing metals from the site of infection, thereby limiting the growth and survival of invading microbes. Critical to this metal-based defense strategy, called nutritional immunity, are immune effectors, including lactoferrin, transferrin and calprotectin, which bind essential metals^1–3^. Further challenging pathogens, the host also actively harnesses the inherent toxicity of metals to disrupt cellular function and inhibit growth^4,5^. As a result, Mn and Fe are not depleted uniformly throughout the body and the bioavailability of metal varies extensively across host tissues^6,7^. Thus, as pathogens disseminate, they must strictly maintain metal homeostasis despite wide fluctuations in metal bioavailability. The essentiality and toxicity of metals are dependent on the environment, including the abundance of other metals^8–10^. Several studies notably highlight close interactions between Fe and Mn homeostasis^11,12^. This indicates a need for bacteria to integrate the availability of both Fe and Mn into the regulatory mechanisms that manage their uptake and efflux. However, the mechanisms that enable this critical integration are unknown.

In response to the manipulation of metal abundance by the immune response, pathogens utilize transcriptional regulators, which directly interact with the cognate metal ion to control adaptive responses. The ferric uptake regulator Fur and the Mn-responsive transcriptional repressor MntR are two well-characterized examples^13,14^. In addition to directly regulating genes involved in metal transport and storage, metal-dependent transcription factors often coordinate a small regulatory RNA (sRNA)-dependent response^15,16^. Small regulatory RNAs, also named trans-acting sRNAs, are usually short RNA molecules, typically ranging from 50 to 500 nucleotides in length^17,18^. They often interact with mRNA targets through imperfect base pairings, altering translation initiation and/or mRNA stability. sRNAs are involved in bacterial adaptation to environment fluctuations, stress responses, virulence, and many other physiological processes. RyhB is the first trans-acting sRNA described as involved in metal-sparing response in *Escherichia coli*^19^. RyhB is negatively regulated by Fur, which directly responds to intracellular Fe levels. Upon Fe starvation, RyhB controls a large set of mRNAs to globally increase Fe uptake, reduce cellular demands for iron, and reallocate Fe to essential biological processes. Over the past two decades, several RyhB-like sRNAs have been discovered among bacteria like PrrF1/2 in *Pseudomonas aeruginosa*^20^, FsrA in *Bacillus subtilis*^21^, and IsrR in *Staphylococcus aureus*^22^. IsrR sRNA is necessary for staphylococcal infection and prevents the expression of iron-demanding enzymes such as those involved in the TCA cycle^23–25^. While studies of metal sparing RNAs have traditionally focused on Fe homeostasis, it is now evident that sRNAs are also crucial in regulating other metallosystems^15^. A perfect example is the Mn-responsive sRNA RsaC in *S. aureus*^26^. RsaC is co-transcribed with the main Mn ABC transporter MntABC and then released by a specific RNase cleavage. Both MntABC and RsaC are produced in response to Mn starvation when the control by MntR is released. RsaC sRNA notably inhibits the synthesis of the Mn-dependent superoxide dismutase A (SodA), promoting the transition to the iron-associated SodM^27^ to reestablish reactive oxygen species (ROS) detoxification pathway in Mn-starved cells.

In addition to trans-acting sRNAs, riboswitches can also play a role in metal homeostasis. Over 50 distinct classes of natural riboswitches, also named cis-acting RNA elements, have been identified so far^28^. These regulatory motifs, embedded in the 5’ untranslated region (5’UTR) of mRNAs, sense and respond to diverse molecules and metabolites including enzyme cofactors, amino acids and sugars. Others respond to metal availability, such as the *yybP-ykoY* riboswitch, which specifically senses intracellular Mn^2+^ levels^29,30^. When riboswitches bind to their cognate ligands, they can regulate downstream gene(s) expression by controlling either the formation of an intrinsic transcription terminator or the sequestration of the ribosome binding site. The *yybP-ykoY* motif was originally discovered in *B. subtilis*, upstream of *yybP* and *ykoY* genes^31^, but it has since been identified in over 1,000 bacterial genomes^29,32^. The crystal structures of the *yybP-ykoY* riboswitch from several organisms have been determined, revealing key roles of Mn^2+^ in the stabilization of the conserved RNA tertiary folding^30,33^. The riboswitch *yybP*-*ykoY* is often located upstream of genes encoding Mn exporters or Mn tolerance proteins such as MntP and Alx in *E. coli*^29,34^, and YybP and YkoY (MeeY) in *B. subtilis*^35,36^. These riboswitch-controlled membrane proteins belong to four major protein families: TerC, P-type ATPase, UPF0016 and MntP^32^. Not all Mn exporters are under the control of the *yybP-ykoY* riboswitch. For example, the cation diffusion facilitator proteins MntE in *S. aureus* and MneP/S in *B. subtilis*^37^ are exclusively controlled by the transcription factor MntR^11^. Remarkably, MntP is regulated by both *yybP-ykoY* riboswitch and MntR in *E. coli*^34^.

In this study, we showed that the sole *yybP-ykoY* (*mnrS*) motif in *S. aureus* controls the synthesis of the Mn efflux pump MntY in response to intracellular Mn levels. Remarkably, we revealed an unprecedented interaction between a trans-acting and a cis-acting regulatory RNA in bacteria. By binding directly to the Mn-sensing riboswitch *mnrS*, the iron-responsive IsrR sRNA alters the conformation of the riboswitch and, consequently, the synthesis of MntY. Further investigation revealed that the interaction between a Fe-responsive sRNAs and a Mn-sensing riboswitch is conserved in multiple bacteria, including *B. subtilis* where FsrA sRNA can interact with both *yybP-ykoY* riboswitches. This unexpected and highly controlled regulation prompted us to investigate the functions of *S. aureus* MntY. We demonstrated that MntY critically contributes to *S. aureus* survival in both low and high Mn environments. In the presence of toxic levels of Mn, MntY facilitates Mn detoxification. In Mn-limiting conditions, the importance of MntY stems from its role in metalating extracellular Mn-dependent proteins such as LtaS, the lipoteichoic acid synthase. Loss of MntY also impaired the ability of *S. aureus* to elaborate virulence factors, resist the antimicrobial activity of calprotectin, survive in the presence of immune cells, and cause infection. The drastic effects observed when its function is altered, the need to precisely regulate its activity and its localization at the membrane suggest that MntY is a promising drug target for staphylococcal infections.

## RESULTS

### IsrR sRNA directly pairs with the Mn-responsive *yybP-ykoY* riboswitch

Although the predicted targetome of IsrR sRNA is broadly similar to those described for RyhB-like sRNAs, one candidate, HG001_00869, stood out in our CopraRNA analysis (Table S1). Notably, this putative target was not evidenced by prior studies^22,25,38^. IsrR is predicted to base pair with nucleotides −184 to −163 from the start codon of HG001_00869. HG001_00869 protein is closely related to YkoY (MeeY) (55% identity) and its paralog YceF (MeeF) in *B. subtilis* (Figures S1A-B). YkoY is, a TerC family membrane protein involved in the efflux and metalation of extracellular Mn-dependent proteins^36^. The structure of YkoY and HG001_00869 predicted using AlphaFold^39^ are also similar (Figure S1C). We will use MntY (<u>Mn t</u>ransporter <u>Y</u>) to designate HG001_00869 hereafter.

Remarkably, the putative IsrR binding site located upstream of *mntY* gene corresponds to the well conserved and widely distributed Mn-responsive *yybP-ykoY* riboswitch (RF00080). The secondary structure of the *S. aureus yybP-ykoY* (Figure 1A), renamed *mnrS* for <u>Mn</u>-responsive riboswitch in *<u>S</u>. aureus* for the sake of simplicity, was inferred from PbAc probing assays (Figure S1D) and previously determined riboswitch conformations in *B. subtilis*, *L. lactis* and *E. coli*^29,30^. In its OFF state, the transcription of *mnrS*-*mntY* is prematurely stopped due to the presence of a Rho-independent terminator (from nts +113 to +147; Figure 1A). The presence of Mn induces conformational changes to prevent premature termination, allowing the formation of the full-length transcript. Based on the crystal structures of several *yybP-ykoY* riboswitches^30,33^, the two conserved L1 and L3 loops form a unique Mn²⁺ binding site that stabilizes the overall folding of the riboswitch (Figures 1A and S1E). This Mn²⁺-dependent structural configuration is essential for regulating the OFF/ON switch, allowing precise control of downstream gene expression.

**Figure 1.**
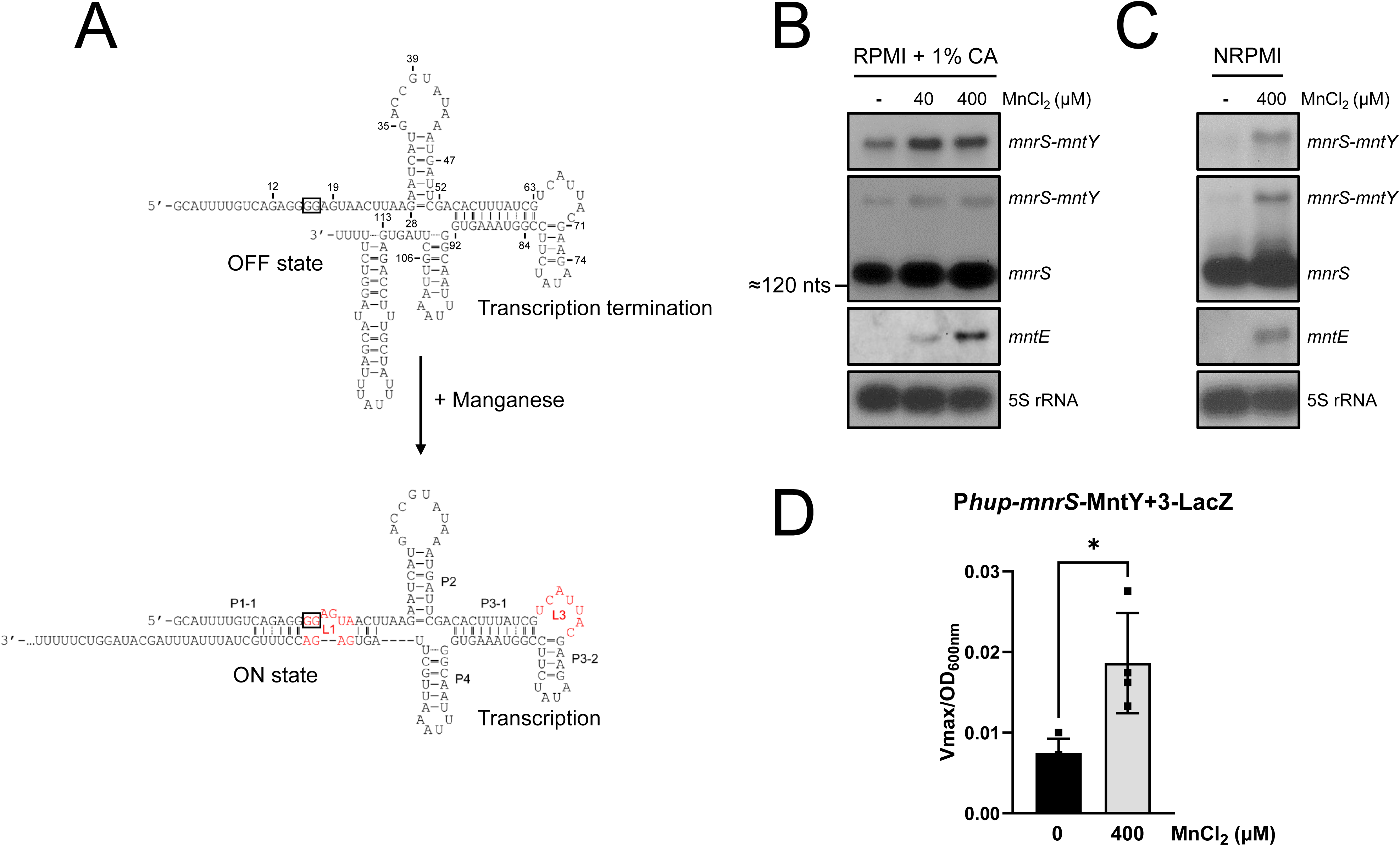
The *mnrS*-*mntY* transcript is Mn-responsive in *S. aureus*. A. Secondary structure of the *S. aureus yybP-ykoY* riboswitch and manganese-induced structural rearrangements inferred from lead acetate (PbAc) probing assays (Figure S1C) and previously characterized riboswitch structures. The GG motif crucial for Mn responsiveness is framed. The highly conserved L1 and L3 loops, which are part of the Mn-binding site, are indicated in red. The +1 of *mnrS* was determined by RNA sequencing^67^. B-C. Northern blot analysis of *mnrS*, *mnrS-mntY* and *mntE* mRNA levels in (B) RPMI supplemented with 1% casamino acids (CA) and (C) chelex-treated NRPMI medium only supplemented with 1 mM MgCl_2_ and 100 µM CaCl_2_. When indicated, 40 or 400 µM of MnCl_2_ were added at the very beginning of culture growth. Two RNA probes targeting either *mnrS* or *mntY (HG001_00869)* sequences were used to distinguish the premature termination from the full-length transcript formation as indicated on the right. 5S rRNA was used as loading and length (≈120 nts) control. Results are representative of at least two independent experiments. D. β-galactosidase assays using *mnrS*-MntY+3-LacZ translational fusion under the control of a constitutive hup promoter (P*hup*) in the absence (dark) or presence of 400 µM MnCl_2_ (gray). Cells were grown in NRPMI medium supplemented with 1 mM MgCl_2_, 100 µM CaCl_2_, 25 µM ZnCl_2_ ± 400 µM MnCl_2_ and harvested at OD_600nm_ = 1. Results are representative of four independent experiments ± SD. A Mann-Whittney test was performed using Prism software (*, P-value<0,05). See also Figure S1.

To facilitate further investigation, the Mn responsiveness of the *mnrS* riboswitch was first confirmed in *S. aureus* (Figure 1). Using Northern blot analysis, a large band corresponding to the full-length *mnrS*-*mntY* transcript was observed using probes targeting either the riboswitch (*mnrS*) or the coding sequence of *mntY* (Figure 1B). The *mnrS* probe also revealed the presence of an abundant ≈150-nt long transcript, suggesting that the *mnrS*-*mntY* locus also produces a terminated riboswitch. The addition of supplemental Mn increased the level of the full-length *mnrS-mntY* transcript, indicating that the abundance of this transcript is dependent on cellular Mn concentrations. The transcript coding for the Mn efflux pump MntE was included as its production is controlled by MntR at the transcriptional level in response to Mn excess^11^. While the level of both mRNAs increases in the presence of high Mn concentrations (40 and 400 µM), the full-length *mnrS-mntY* mRNA is present even in the absence of supplemental Mn (Figure 1B). The *mnrS-mntY* transcript was only drastically reduced in a chelex-treated NRPMI medium, where Mn is depleted (Figure 1C). This suggests that the presence of Mn affects the production of MntY protein via the formation of an anti-terminator structure, corresponding to the ON state of the *mnrS* riboswitch. This hypothesis was tested with a translational *lacZ* fusion using β-galactosidase assays. The endogenous *mnrS-mntY* promoter was replaced by the constitutive promoter of the *hup* gene to eliminate any potential MntR transcriptional regulation (Figure S1F). We observed that the activity of the MntY+3-LacZ fusion increases in the presence of 400 µM (2.5-fold), demonstrating that the *mnrS* riboswitch is responsible for this Mn-dependent activation (Figure 1D). As mentioned above, *mnrS-mntY* transcription is not completely abolished at low or physiological Mn levels. Bastet et al.^40^ reported that the premature termination efficiency of *mnrS-mntY* transcript is affected by a mispairing in the riboswitch terminator (Figure S1G). However, the read-through observed in Figures 1B-C cannot be solely attributed to terminator leakage as the mutated terminator (U>A; more stable stem) did not lower the level of full-length *mnrS-mntY* mRNA (Figure S1H). This result sheds light on the conformational flexibility of the *mnrS* riboswitch at low Mn levels.

To test the in silico prediction showing that IsrR sRNA binds to *mnrS* riboswitch, we performed electrophoresis mobility shift assays (EMSA; Figure 2A) using in vitro transcribed RNAs. We used either the 148-nt long *mnrS* transcript or a longer transcript ending at nucleotide +115 in the coding sequence of *mntY* (referred as *mnrS*-*mntY*). We showed that the 5’ end radiolabeled IsrR forms a high-affinity complex with both *mnrS* or *mnrS*-*mntY* in vitro, with a dissociation constant (Kd) of approximatively 17 nM and 15 nM, respectively (Figure S2A). We obtained similar result by performing the opposite experiment with the 5’ end of the *mnrS* riboswitch radiolabeled and cold IsrR (Figure S2B). Altogether, this suggests that IsrR binds to the riboswitch rather than the ribosome binding site or the coding sequence of *mntY*. As the *mnrS* riboswitch is subjected to conformational changes in the presence of Mn^2+^ ions^29,30^, we evaluated the impact of 10 mM MnCl_2_ on the complex formation (Figures S2C-D). Even in the presence of Mn, IsrR binds efficiently to *mnrS* in vitro. Terminator efficiency assays, in which the promoter P*lysC* was fused to the 5’UTR of *mntY*, including *mnrS* riboswitch, confirmed that 1 mM MnCl_2_ is sufficient to induce the ON state of *mnrS* riboswitch in vitro (Figure S2E). These results suggests that IsrR can interact with the *mnrS* riboswitch regardless of its secondary structure rearrangement in the presence or absence of its ligand.

**Figure 2.**
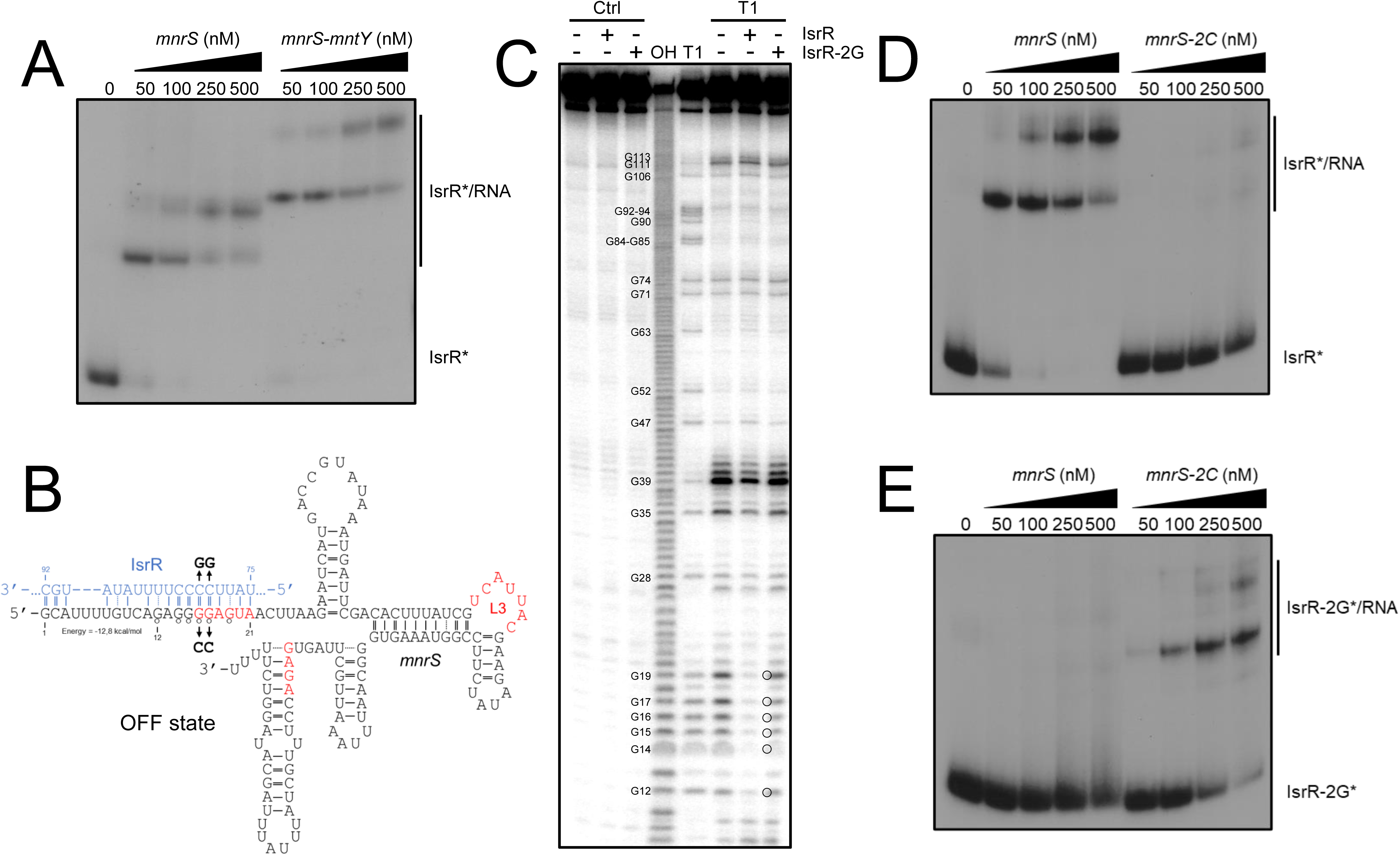
IsrR sRNA directly binds to *mnrS* riboswitch in vitro. A. Gel retardation assays using IsrR sRNA (full-length), *mnrS* riboswitch (full-length) and *mnrS*-*mntY* mRNA (from −183 to +115). 5’ end-radiolabeled IsrR (*) was incubated with increasing concentrations of *mnrS* or *mnrS*-*mntY* (0, 50, 100, 250 and 500 nM). B. The IsrR-*mnrS* pairing site was in silico predicted using IntaRNA algorithm^44^. The mutation of pairing sites (*mnrS*-2C and IsrR-2G) are indicated by arrows. Nucleotides whose levels decrease in presence of IsrR in Figure 2C are indicated with open circles. IsrR is shown in blue. Sequences involved in the formation of the L1 and L3 loops are indicated in red. C. RNase T1 probing of 5’end-radiolabeled *mnrS* riboswitch (full-length) incubated in the absence (-) or presence of IsrR or IsrR-2G. Ctrl, non-reacted controls; OH, alkaline ladder; T1, RNase T1 ladder. The numbers to the left show guanine (G) positions with respect to the transcription start (+1) of *mnrS-mntY*. IsrR-induced decreased cleavages at G residues are indicated with open circles. Gel retardation assays using either (D) IsrR or (E) IsrR-2G and increasing concentrations of *mnrS* or *mnrS*-2C (from 0 to 500 nM). Results are representative of at least two independent experiments. See also Figure S2.

According to in silico predictions, IsrR targets a G-rich motif at the beginning of the *mnrS* riboswitch sequence (Figure 2B). This region is highly conserved among bacteria and was previously described as crucial for Mn responsiveness^29^. Indeed, mutations in the G-rich motif alter the formation of the metal binding site and hinder the Mn-dependent induction (Figure S1E). To validate this pairing site, we performed in vitro RNase T1 protection assays using 5’end-radiolabeled *mnrS* riboswitch. As expected, all the guanines from nts 12 to 20 were cleaved, indicating that they are located in a single-stranded region in the absence of Mn (Figure 2C). In the presence of IsrR, this region become fully protected from RNase T1 cleavage, supporting the formation of intermolecular base pairings. Mutation of the putative IsrR base pairing site (C79G and C80G; IsrR-2G) hindered binding to *mnrS*, as protection from RNase T1 cleavage was lost in the presence of IsrR-2G. Consistent with these data, a mutation in the *mnrS* sequence (G16C and G17C; *mnrS*-2C) prevented IsrR-*mnrS* interaction (Figure 2D), whereas the complex formation was restored using the compensatory mutants *mnrS*-2C and IsrR-2G (Figure 2E). Taken together, these results validate a direct interaction between IsrR and *mnrS* riboswitch, involving a critical region required for switching to the ON state (Figure 2B).

### IsrR sRNA directly impairs the synthesis of full-length *mnrS*-*mntY*

Since IsrR interacts with the nucleotides involved in the anti-terminator structure of the *mnrS* riboswitch, which favors the OFF state, we hypothesized that it leads to premature termination of *mnrS-mntY* transcription. To test this, we monitored the level of *mnrS* and *mnrS*-*mntY* transcripts in WT and Δ*isrR* mutant strains by Northern blot using probes targeting either the *mnrS* riboswitch or the coding sequence of *mntY* (Figure 3A). A significant increase in the full-length mRNA level (2.45-fold) was detected in the absence of IsrR with both probes (Figure 3B), suggesting that IsrR lowers the synthesis of the full-length mRNA. To determine whether this is due to the direct interaction between IsrR and *mnrS* riboswitch as demonstrated in vitro (Figure 2), we monitored the level of the full-length mRNA in an *isrR*-2G mutant background. The *isrR-2G* mutation results in a similar increase in the full-length *mnrS-mntY* mRNA level as the *isrR* deletion (Figure S3A). These results indicate that IsrR interacts directly with the *mnrS* riboswitch to reduce the full-length transcript levels in vivo.

**Figure 3.**
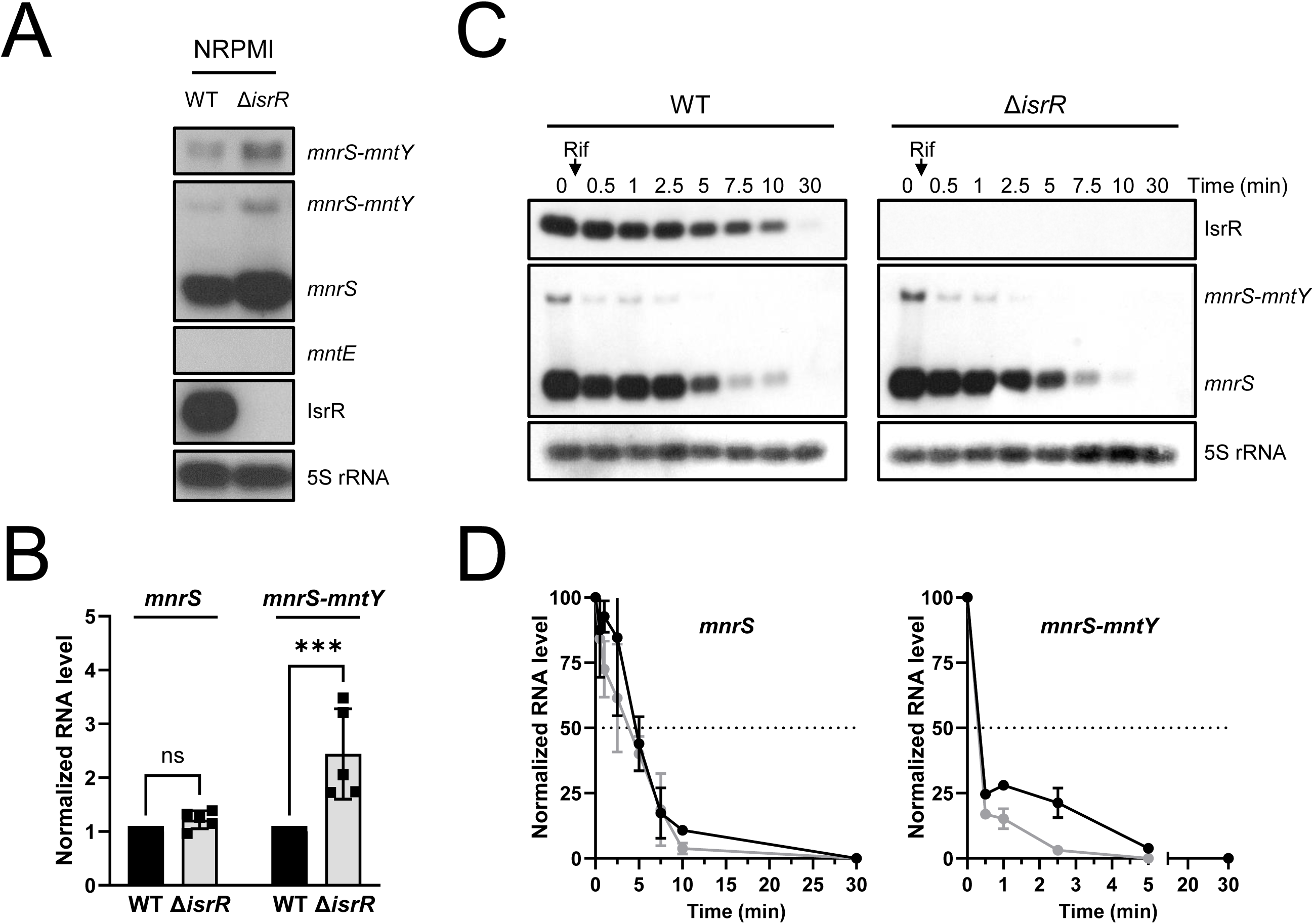
IsrR alleviates production of the full-length *mnrS-mntY* transcript by inducing premature termination in response to Fe starvation. A. Northern blot analysis of *mnrS* and *mnrS-mntY* RNA levels in WT and Δ*isrR* backgrounds in NRPMI medium only supplemented with 1 mM MgCl_2_ and 100 µM CaCl_2_. Cells were harvested at OD_600nm_=1. The *mntE* mRNA and 5S rRNA were used as negative control and loading control, respectively. Results are representative of five independent experiments. B. Densitometric analysis of Figure 3A performed using ImageJ software (n=5). RNA levels were normalized by 5S rRNA and relativized to the WT condition. A two-way ANOVA analysis followed by Sidak’s multiple comparison test was performed using Prism software (***, P-value<0.0005; ns, non-significant). C. Determination of *mnrS* and *mnrS-mntY* mRNA half-life using rifampicin assays in WT and Δ*isrR* strains. IsrR expression was triggered by 2,2-dipyridyl (250 μM; DIP) at OD_600nm_=1. After 15 min, the transcription was block by addition of rifampicin (300 µg/mL). Total RNA was then extracted at indicated times. The 5S rRNA is used as a loading control. Results are representative of three independent experiments. D. Densitometric analyses of Figure 3C performed using ImageJ software (n=3). RNA levels were normalized by 5S rRNA and relativized to respective T0. See also Figure S3.

In the Northern blots, we occasionally observed a slight but non-significant increase in the band corresponding to the *mnrS* riboswitch in the Δ*isrR* background (Figures 3A-B), raising the possibility that the *mnrS*-IsrR interaction could trigger mRNA decay. Thus, we tested the stability of the *mnrS* riboswitch and *mnrS*-*mntY* mRNA using rifampicin assays in wild type and Δ*isrR* mutant strains (Figure 3C). Consistent with our previous observations (Figures 3A-B), the level of the full-length mRNA was higher at t0 in absence of IsrR (Figure 3C). However, there was no difference in the *mnrS*-*mntY* (t_1/2_ ≈ 20 sec) or *mnrS* riboswitch (t_1/2_ ≈ 5 min) half-life in WT and Δ*isrR* backgrounds (Figure 3D). This indicates that IsrR does not impact the stability of either the full-length mRNA or the riboswitch. Combined with our prior results, this indicates that IsrR reduces the synthesis of full-length *mnrS-mntY* mRNA by inducing a conformational change of the *mnrS* riboswitch, triggering premature termination at the co-transcriptional level.

Having observed an interaction between IsrR and *mnrS*, the possibility of an interaction with the other staphylococcal efflux pump MntE was explored. However, no clear interaction between IsrR and *mntE* mRNA was observed in vitro using EMSA (Figure S3B). This suggests that, unlike MntE, Fe abundance is directly integrated into the regulatory scheme that controls MntY synthesis.

### The *mnrS*-IsrR interaction is widely conserved in Firmicutes

Both *yybP-ykoY* riboswitches (Figure S4A) and RyhB-like sRNAs are widespread among bacteria^16,29^. This suggests that the interaction between the iron-sparing response regulator and the Mn-dependent riboswitches could be conserved. The model organisms *E. coli* and *B. subtilis* both possess a RyhB analog (RyhB and FsrA, respectively) and two copies of the *yybP-ykoY* riboswitch. Using multiple sequence alignment, we noticed that the pairing site on the riboswitch sequence is highly conserved (Figure S4B).

To go further, we studied the co-appearance and interaction potential of Fe-responsive sRNAs and Mn-sensing *yybP-ykoY* riboswitches in bacteria (Figure 4A). We compiled >8500 *yybP-ykoY* riboswitch homologs and categorized them based on their downstream gene (Table S2). The vast majority of the riboswitches belong to three families (2671 for MntP, 2429 for YkoY and 1647 for Gdt1/MneA). In parallel, we scanned for homologs of the Fe stress responsive sRNAs ArrF^41^, FsrA^21^, Isar1^42^, IsrR^22^, NrrF^43^, PrrF1/2^20^ and RyhB^19^. The interaction potential of the sRNAs with the respective riboswitch in each organism was investigated with IntaRNA^44^ (Table S2). A minimal interaction energy of ≤-9 kcal/mol and an interaction position covering the 5’ “GGGGAGUA” motif in the riboswitch sequence were chosen as criteria for a potential functional interaction based on the EMSA data (Figures 4D-F and S4C). Given the above assumptions, only the YkoY and YybP riboswitch families showed a strong interaction potential with IsrR and FsrA sRNAs (Figures 4A-C).

**Figure 4.**
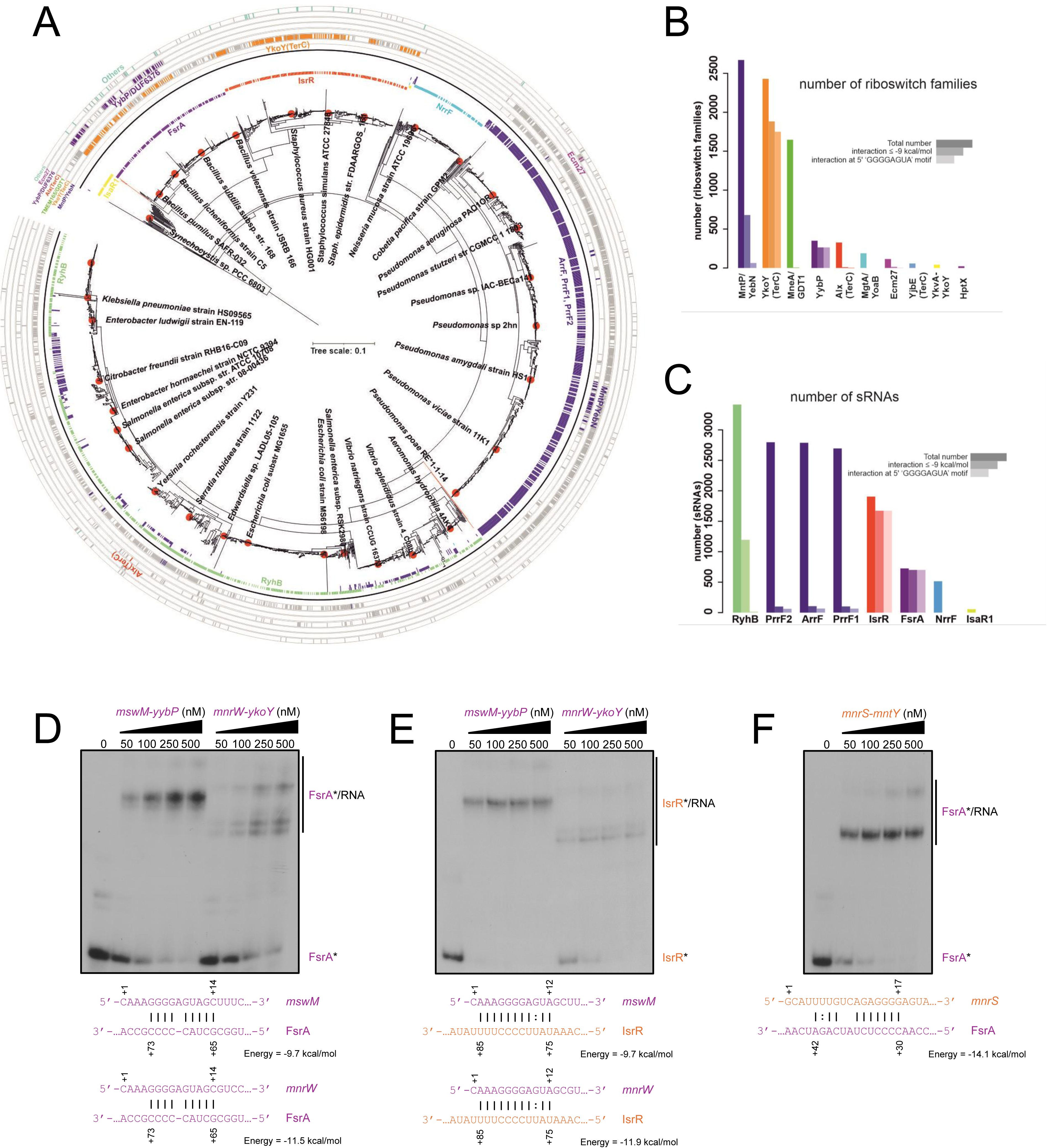
The interaction between *yybP-ykoY* riboswitches and iron-responsive sRNAs is conserved across bacteria. A. Appearance and interaction-potential of *yybP-ykoY* riboswitch families and Fe stress related sRNAs. The graph represents the 16S rDNA-based NJ tree of riboswitch and sRNA (co)-appearance. The outer ring shows the occurrence patterns of the 9 most frequent riboswitch families as bars. Colors (one per family) indicate the predicted interaction potential of each riboswitch with one of the sRNAs shown in the inner ring. The gray color indicates an interaction energy >9 kcal/mol or no predicted 5’GGGGAGUA-site. Closely related organisms were condensed to one representative based on the 16S rDNA similarity. B. Numbers of detected homologs for the 11 most frequent riboswitch families (left bars = total numbers; middle bar = predicted interaction with energy ≤-9 kcal/mol; right bar = predicted energy ≤-9 kcal/mol & interaction at 5’ end covering the 5’-GGGGAGUA motif). The colors correspond to the colors used in (A). C. Numbers of detected homologs for the 8 investigated Fe stress related sRNAs. See (B) for more details. D. Gel retardation assays using FsrA (full-length), *mswM-yybP* (from −307 to +172) and *mnrW-ykoY* (from −21 to +289) transcripts from *B. subtilis* 168. 5’ end-radiolabeled FsrA sRNA (*) was incubated with increasing concentrations of cold abovementioned mRNAs (0, 50, 100, 250 and 500 nM). Putative pairing sites predicted in silico using IntaRNA algorithm are indicated below. Results are representative of at least two independent experiments. E. Gel retardation assays using IsrR (full-length), *mswM-yybP* (from − 307 to +172) and *mnrW-ykoY* (from −21 to +289) transcripts. RNAs from *S. aureus* and *B. subtilis* are shown in orange and purple, respectively. G. Gel retardation assays using FsrA sRNA (full-length) and *mnrS*-*mntY* mRNA (from −183 to +115). See also Figure S4.

Thus, the sRNA-riboswitch interaction seems to be conserved in *B. subtilis* but not in *E. coli*. To validate our in silico analysis, we performed EMSA to determine if RyhB and FsrA can interact with their respective *yybP-ykoY* riboswitches in vitro. While FsrA efficiently binds to both *mswM-yybP* and *mnrW-ykoY* transcripts (Figure 4D), RyhB does not interact with *yybP-ykoY-mntP* and *sraF-alx* transcripts (Figure S4C). The hybridization energies involving RyhB are quite unfavorable compared to those predicted for *S. aureus* and *B. subtilis* (Figures 2B and 4D). In addition, we used translational fusions to measure the potential impact of RyhB in vivo. The native promoter was replaced by Plac to avoid any regulation dependent on MntR. In agreement with the EMSA, the overexpression of RyhB does not impact the β-galactosidase activity of the translational Plac-*yybP-ykoY*-MntP-LacZ and Plac-*sraF*-Alx-LacZ fusions in *E. coli* (Figure S4D-E). As expected, each construct was activated in the presence of Mn.

Finally, we analyzed the formation of RNA duplexes by mixing a RyhB-like sRNA from one species with a riboswitch from another. We noticed that IsrR sRNA from *S. aureus* strongly binds to both *mswM-yybP* and *mnrW-ykoY* transcripts from *B. subtilis* (Figure 4E). Similarly, FsrA sRNA from *B. subtilis* pairs with *mnrS-mntY* transcript from *S. aureus* (Figure 4F). This cross-species interaction provides further evidence that this sRNA-riboswitch interaction is conserved across Firmicutes, although there is no apparent homology between IsrR and FsrA.

### The deletion of *mnrS-mntY* induces strong phenotypic defects

The homology of MntY with known Mn efflux pumps suggests that it contributes to *S. aureus* Mn homeostasis. As a first step to test this hypothesis, a Δ*mnrS*-*mntY* mutant was created and examined. Since MntY is a predicted Mn efflux pump, a Δ*mntE* mutant was evaluated in parallel. Surprisingly, during routine cultivation on blood agar plates the Δ*mnrS*-*mntY* mutant, but not the Δ*mntE* mutant, differed from WT (Figure 5A). The Δ*mnrS-mntY* colonies were tiny and cream colored, having lost the characteristic golden color of *S. aureus* and lacking hemolysis. Similar results were observed when the Δ*mnrS*-*mntY* mutant was plated on BHI agar (Figure S5A). Microscopic examination revealed no gross differences in the size and shape of the bacterial cells (Figure S5B). The Δ*mnrS-mntY* mutant also lacked proteolytic activity on 5% skim milk plates (Figure S5C) when compared to the WT strain. Finally, deletion of the *mnrS-mntY* locus caused hypersensitivity to daptomycin (Figure S5D). A Δ*mnrS* mutant did not impact any of the assayed phenotypes, while expression of either *mnrS-mntY* or *mntY* under the control of a weak promoter (pEW plasmid^45^) complemented the defects of the Δ*mnrS*-*mntY* mutant, including hemolysis activity (Figure 5A).

**Figure 5.**
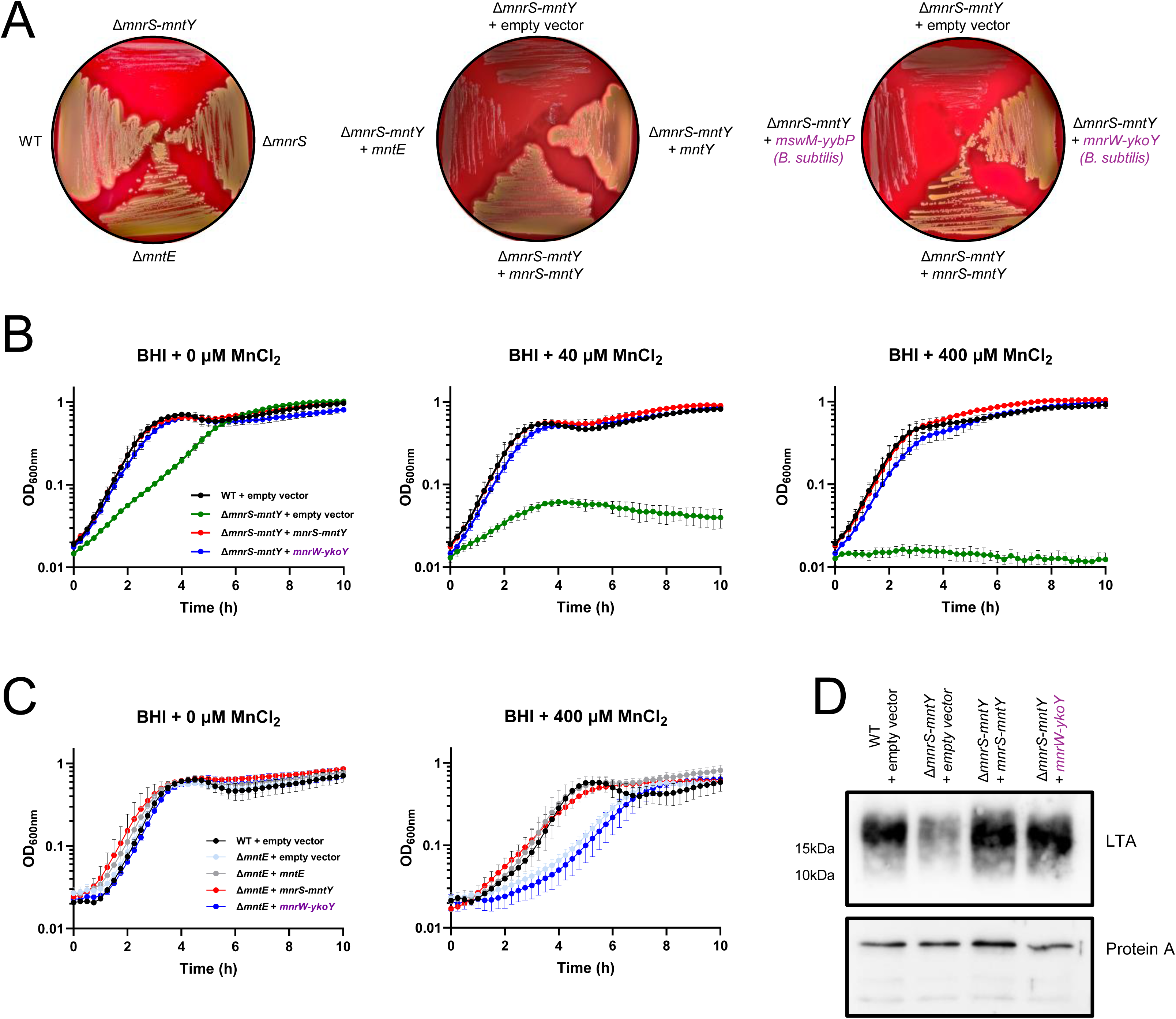
*mntY* is required for Mn detoxification and LTA synthesis in *S. aureus*. A. *S. aureus* strains were streaked onto blood agar plates and incubated at 37°C for 48h. The Δ*mnrS-mntY* mutant strain was complemented with a pEW derivate plasmid which enables the constitutive expression of indicated genes (*mntY*, *mnrS-mntY*, *mntE, mnrW-ykoY* or *mswM-yybP*). Gene sequences from *Bacillus subtilis* are shown in purple. The empty vector pEW was used as control. Blood agar plates were supplemented with erythromycin (10 μg/mL) for plasmid selection. B. Growth monitoring (OD_600nm_) in BHI medium supplemented with 0, 40 or 400 μM MnCl_2_ and 10 μg/mL erythromycin for 10h at 37°C. WT and Δ*mnrS-mntY* strains contain the empty vector or a pEW derivate plasmid which enables the constitutive expression of *mnrS-mntY* or *mnrW-ykoY* from *B. subtilis*. Data correspond to the mean of three independent experiments ± standard deviation (SD). C. Growth monitoring carried out under the same conditions as (B). WT and Δ*mntE* strains contain the empty vector or a pEW derivate plasmid which enables the constitutive expression of *mntE*, *mnrS-mntY* or *mnrW-ykoY* from *B. subtilis*. D. Immunodetection of lipoteichoic acid (LTA) using cell extracts from WT+pEW, Δ*mnrS- mntY*+pEW, Δ*mnrS-mntY*+pEW-*mnrS-mntY and* Δ*mnrS-mntY*+pEW-*mnrW-ykoY* from *B. subtilis*. Bacteria were grown to mid exponential growth phase in BHI media (OD_600nm_ ≈ 1). Equal amounts of whole-cell lysates were separated by 20% SDS-PAGE, transferred to PVDF membranes and immunodetected with anti-LTA monoclonal antibodies. Protein A was used as a loading control. Results are representative of two independent experiments. See also Figure S5.

As the Δ*mnrS*-*mntY* mutant had a profound growth defect, the media was changed to BHI (Figure 5B). The addition of Mn to the medium decreased the ability of Δ*mnrS*-*mntY* to grow consistent with a role in Mn detoxification. Unexpectedly, the removal of Mn from the medium via chelex treatment did not fully rescue the growth of the Δ*mnrS*-*mntY* mutant (Figure S5E), indicating that its role is not limited to Mn tolerance. Notably, the expression of *mntY* under the control of a strong promoter (pES plasmid^46^) appears toxic as strains expressing *mntY* from the pES plasmid could not be recovered. However, a pES plasmid containing *mnrS*-*mntY* could be introduced into *S. aureus*. Cumulatively, these observations reveal that loss of MntY has a profound impact on *S. aureus*, but that its expression must be tightly controlled, with the *mnrS* riboswitch critically contributing to this task.

Given the differences in expression between *mntY* and *mntE* (Figures 1B-C), it is not surprising that loss of MntY does not phenocopy loss of MntE. However, similar to the Δ*mntE* mutant (Figure 5C), the Δ*mnrS*-*mntY* mutant has a growth defect in the presence of excess Mn (Figure 5B). This leads to the hypothesis that the growth defects of the Δ*mnrS-mntY* mutant are due to intracellular Mn accumulation and toxicity. To test this hypothesis, *mntE* was expressed from a plasmid under the control of a weak constitutive promoter (pEW). The ectopic expression of *mntE* did not complement the phenotypic and growth defects of the Δ*mnrS*-*mntY* mutant (Figures 5A and S5F), but reversed the sensitivity of the Δ*mntE* mutant to Mn toxicity (Figure 5C), indicating that *mntE* was expressed. Similar to *mntY*, overexpression of *mntE* appears toxic to *S. aureus* as colonies carrying *mntE* in the pES plasmid could not be recovered.

A Δ*ykoY* Δ*yceF* double mutant strain in *B. subtilis* displays, amongst other phenotypes, a significant decrease in colony size^36^. Since MntY shares 55% of sequence identity with YkoY, we evaluated if *mnrW-ykoY* could complement the growth defect of the Δ*mnrS-mntY* mutant. We also used *mswM-yybP* to complement the Δ*mnrS-mntY* mutant as the *yybP-ykoY* riboswitch controls the synthesis of both YkoY and YybP in *B. subtilis*. The *mnrW-ykoY* construct, but not *mswM-yybP*, reversed both the phenotypic and growth defects of the Δ*mnrS-mntY* mutant (Figures 5A-B). Cumulatively, these observations indicate that while phenotypes associated with loss of MntY are driven by a failure to secrete Mn, the role of MntY extends beyond the removal of Mn from the cell.

### MntY is involved in Mn detoxification and metalation of Mn-dependent exoenzymes in *S. aureus*

The totality of the current observations leads to the hypothesis that MntY contributes to the ability of *S. aureus* to detoxify Mn, but also has a second critical function. To test the first aspect of this hypothesis, we overexpressed *mnrS-mntY* in an Δ*mntE* mutant (Figure 5C). As expected, the growth of the Δ*mntE* mutant is reduced by high Mn levels. The growth defect of the Δ*mntE* mutant was rescued by either the expression of *mntE* or *mnrS-mntY* construct. In conjunction with the observation that Δ*mnrS-mntY* is sensitive to levels of Mn, this indicates that MntY contributes to protecting *S. aureus* from Mn toxicity. Notably, although the expression of *ykoY* rescued the growth defect of the Δ*mnrS-mntY* mutant, it did not complement the Δ*mntE* mutant. This indicates that despite their homology, MntY and YkoY have non-redundant cellular functions.

Recently, YkoY and YceF were suggested to contribute to the metalation of extracellular Mn-dependent enzymes in *B. subtilis*, including the lipoteichoic acid (LTA) synthase, LtaS^36,47^. This suggests that the second critical function of MntY is the metalation of extracellular Mn-dependent enzymes. Therefore, to test the second aspect of our hypothesis, the production of LTA was assessed via immunoblotting. When compared to wild type, the Δ*mnrS-mntY* mutant has substantially reduced levels of LTA (Figure 5D). As a control, Protein A levels remain unaffected. The expression of *mnrS-mntY* from a plasmid restores LTA levels. Notably, while YkoY did not rescue the ability of the Δ*mntE* mutant to cope with Mn toxicity, it did restore LTA production. This indicates that although both MntY and MntE are involved in Mn detoxification, MntY is only responsible for exoenzyme metalation in *S. aureus*. It also further supports the idea that MntY and YkoY have related but distinct functions, with MntY contributing both to Mn detoxification and to the activity of Mn requiring extracellular enzymes.

### The Mn efflux pump MntY is crucial for *S. aureus* during infection

MntY is important for the management of Mn within *S. aureus* and this metal crucially contributes to the ability of *S. aureus* to evade host defenses. Therefore, we hypothesized that MntY would contribute to the ability of *S. aureus* to overcome nutritional immunity and the antimicrobial activity of immune cells. First, we investigated the ability of the Δ*mnrS*-*mntY* mutant to grow in the presence of calprotectin, which restricts the availability of Mn and other metals from invaders during infection and is present at sites of infection in excess of 1 mg/mL. When compared to WT bacteria, the Δ*mnrS*-*mntY* mutant was significantly more sensitive to calprotectin with a strong growth defect observed at concentrations as low as 120 µg/mL (Figure 6A). Next, the ability Δ*mnrS*-*mntY* to survive phagocytic killing was evaluated. The mutant was 2-fold more sensitive to killing by macrophage like THP-1 cells (Figure 6B). We then assessed the distribution of bacteria between the extracellular and intracellular environment. With the Δ*mnrS*-*mntY* mutant, 59% of the viable bacteria were intracellular compared to the 6% with the WT strain (Figure 6C). We repeated both assays in the presence of human neutrophils (Figures 6D-E). Similar to macrophages-related assays, the Δ*mnrS*-*mntY* mutant is more sensitive to killing by neutrophils and tends to be phagocytized at higher levels than the WT strain.

**Figure 6.**
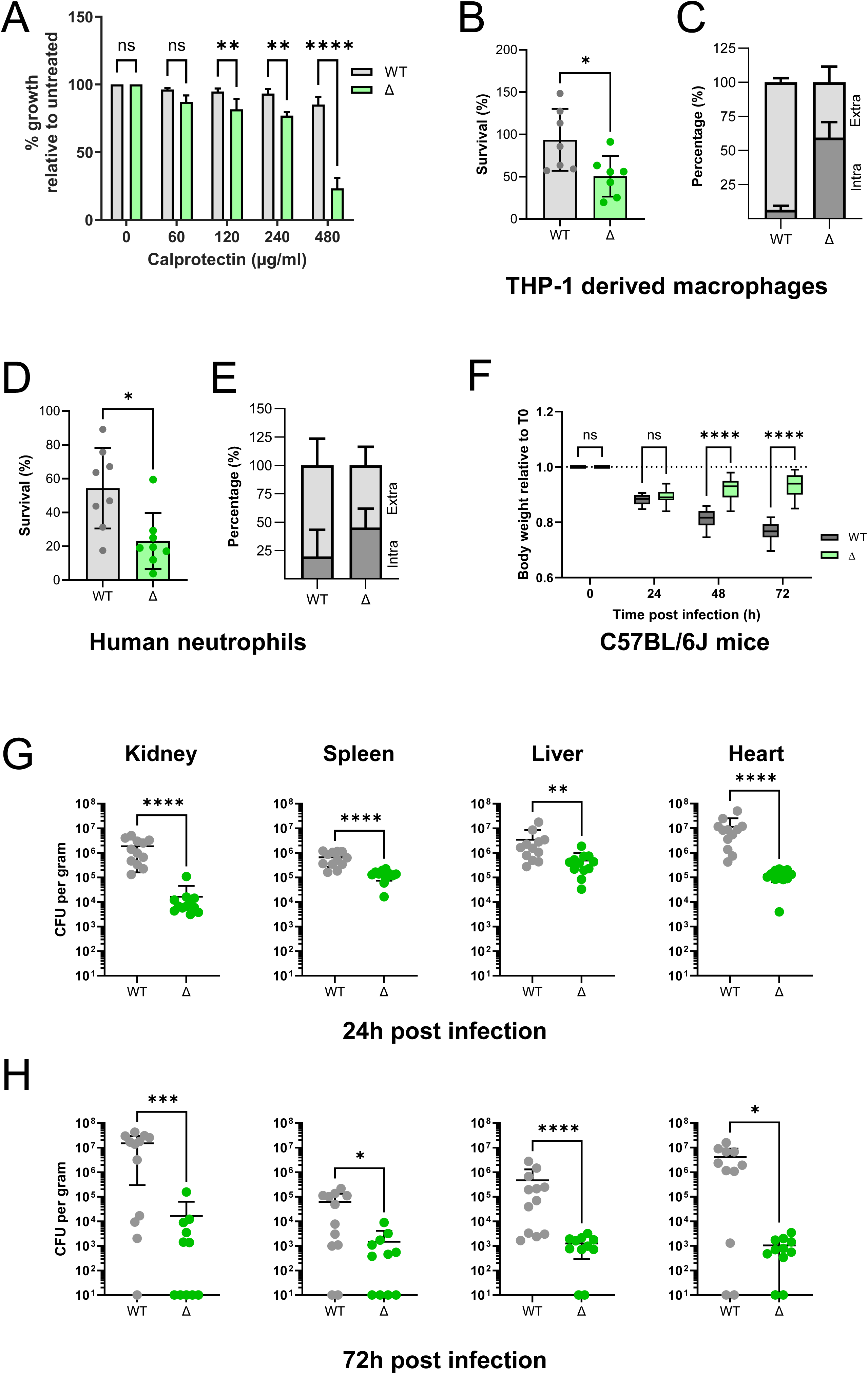
The Mn efflux pump MntY is crucial for *S. aureus* infection. A. Wild type (gray) and Δ*mnrS-mntY* (green) cells were grown in the presence of increasing concentrations of calprotectin (0 to 480 µg/mL). OD_600nm_ was measured after 8h of growth and indicated as percentage of growth relative to untreated. Results are representative of three independent experiments ± SD. A two-way ANOVA analysis followed by Sidak’s multiple comparison test was performed using Prism software (**, P-value<0.005; ****, p-value<0.0001; ns, non-significant). B, D. Survival of wild type (WT; gray) and Δ*mnrS-mntY* (Δ; green) cells assessed after exposure to THP-1-derived macrophages (B; n=7) or human purified neutrophils (D; n=8). Bacterial loads were determined by serial-dilution and plating on BHI agar. The CFU counts were normalized to the initial inoculum (%). C, E. Percentage of extracellular (light gray) and intracellular (dark gray) cells. The bacterial loads were determined for wild type (WT; gray) and Δ*mnrS-mntY* (Δ; green) cells after exposure to immune cells (extracellular) and after internalization and treatment with lysostaphin/gentamicin (intracellular). The CFU counts were normalized to total bacteria (%). F. Mouse weight loss due to infection with WT (dark gray) and Δ*mnrS-mntY* (green) strains. Results were relativized to initial body weight. Results are from two independent experiments (n=12). G-H. Wild type (WT; gray) and Δ*mnrS-mntY* (Δ; green) strains were injected retro-orbitally into C57BL/6J mice. Kidneys, spleen, liver, and heart were collected at 24 (G) and 72 h (H) post-infection, and bacterial loads were determined by serial-dilution and plating on BHI agar. The CFU counts per gram of each organ were reported. Results are from two independent experiments (n=12). A Mann-Whittney test was performed using Prism software (*, P-value<0.05; **, P-value<0.005; ***, P-value<0.0005; ****, P-value<0.0001).

The susceptibility of the Δ*mnrS*-*mntY* mutant to multiple arms of the immune response suggests that it should be important during infection. To test this hypothesis, the virulence of WT and the Δ*mnrS*-*mntY* mutant strains were evaluated using a systemic infection model. Mice infected with the Δ*mnrS*-*mntY* mutant displayed substantially fewer signs of illness and began to recover weight after 48 h of infection, while mice infected with WT strain continued to lose weight for the duration of the experiment (Figure 6F). Bacterial load in kidney, spleen, liver, and heart was also assessed at 24 and 72 h post-infection (Figures 6G-H). Mice infected with Δ*mnrS*-*mntY* mutant had lower bacterial burdens when compared to WT at both time points in all of the tissues assayed.

Altogether, these results demonstrate that the *mnrS-mntY* is critical for *S. aureus* virulence and/or survival during murine and human infection.

## DISCUSSION

Life is constantly faced with the need to adapt to environmental changes. This is particular true for microbes, which have a limited ability to impact their environment. One challenge they face is responding to changes in metal availability. These nutrients are critical for all forms of life but can also be toxic. This duality is acutely faced by pathogens as the host actively restricts the ability of these nutrients and harnesses their toxic properties to combat infection. Leveraging *S. aureus* as a model, the present study revealed a previously unappreciated regulatory node that enables bacteria to coordinate their response to Mn and Fe availability (Figure 7). At this node, the Fe-responsive sRNA IsrR interacts with the Mn-sensing riboswitch *mnrS*. This enables *S. aureus* to control the expression of the Mn efflux pump MntY, which the current investigations revealed is critical for the bacterium to cope with manganese toxicity, maintain essential cellular processes, and cause infection. The integration of the response to Fe scarcity with Mn efflux highlights how microbes must integrate multiple cues when managing metal homeostasis so as to preserve the function of essential processes, while not sensitizing themselves to intoxication. Further investigations revealed that the interaction is conserved within the Firmicutes, highlighting that while the interaction between an sRNA and a riboswitch is unprecedented, the mechanism is widely used to control gene expression and integral to the ability of bacteria to respond to their environment.

**Figure 7.**
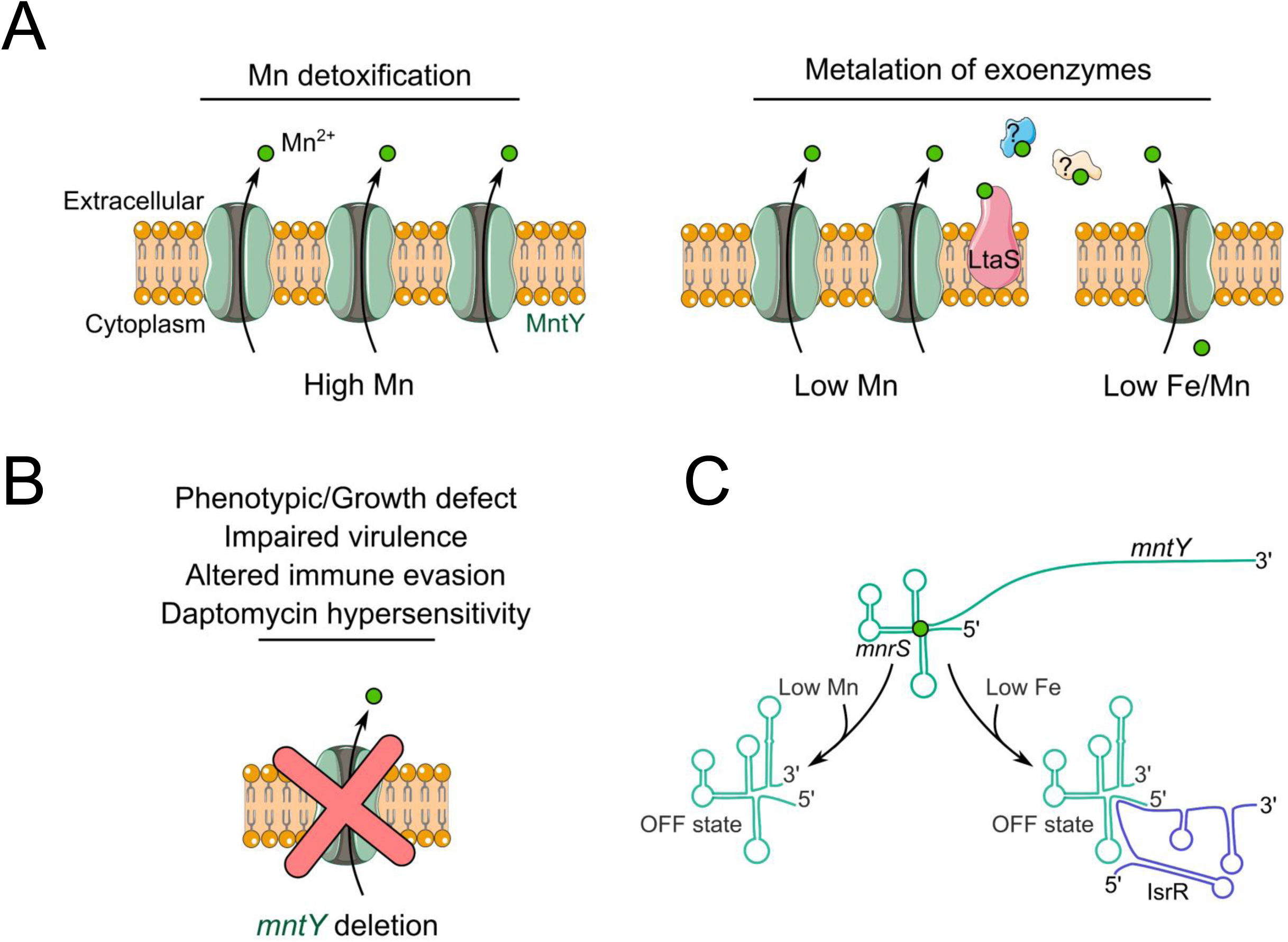
Schematic representation of Mn efflux pump MntY function and regulation. A. The Mn efflux pump MntY is vital for the adaptation of *S. aureus* to both low and high Mn environments, due to its dual role in Mn detoxification and metalation of Mn-dependent exoenzymes, such as LtaS. B. Loss of MntY causes drastic phenotypic and growth defects, impairs virulence, immune evasion and antibiotic resistance. C. The expression of *mntY* is fine-tuned by an unprecedented interaction between a Mn-sensing riboswitch mnrS and a Fe-responsive sRNA IsrR.

### An unprecedented interaction between a cis- and a trans-acting RNA

Bacteria employ both cis-acting (i.e., riboswitches) and trans-acting RNAs (referred to here as sRNAs) to finely regulate gene expression. The synthesis of the membrane transporter BtuB in *E. coli* was recently shown to be controlled by both an adenosylcobalamin riboswitch and OmrA sRNA via distinct and independent regulatory mechanisms^48^. The current investigations revealed a new regulatory combination, the direct interaction between a cis- acting riboswitch (*mnrS*) and a trans-acting sRNA (IsrR). Both RyhB analogs and Mn-sensing riboswitches (*yybP-ykoY* homologs) are widely distributed in bacteria (Figures 4A and S4A). We observed that the targeted sequence on the *yybP-ykoY* riboswitch is also highly conserved (Figure S4B), being involved in the formation of the Mn^2+^ binding site (Figure S1E). A global RNA-RNA interaction prediction, confirmed by targeted validation analysis, showed that the interaction between an iron-responsive sRNA and a *yybP-ykoY* riboswitch is not restricted to *S. aureus,* and is likely to broadly occur in Firmicutes and some Gram-negative bacteria including *Pseudomonas* species (Figure 4). It is probable that this unprecedented regulatory mechanism extends beyond Mn-sensing riboswitches and Fe-responsive sRNAs, as several omics-based analyses have observed potential riboswitch-sRNA interactions. Examples include a T-box riboswitch and RsaG sRNA in *S. aureus*^49^, and a guanidine-II riboswitch and ArcZ sRNA in *E. coli*^50^.

Trans-acting sRNAs have long been thought to act post-transcriptionally on fully transcribed mRNA. However, recent work revealed that the Hfq-associated DsrA sRNA binds faster to the nascent *rpoS* transcript than to the refolded one in *E. coli*^51^. This led to the suggestion that sRNA may also bind to the nascent mRNA during transcription and prior to folding. Here, we demonstrated that binding of IsrR precludes the formation of the anti-terminator structure of the *yybP-ykoY* riboswitch at the co-transcriptional level, and consequently, induces the premature termination of the *mnrS-mntY* transcript (Figure 3).

The *mnrS-mntY* locus is constitutively expressed, leading the *mnrS* riboswitch to accumulate under normal conditions (Figure 1). Notably, the half-life of the *mnrS* riboswitch (t_1/2_ ≈ 5min) (Figure 3C) is considerably longer that the vast majority of mRNAs^52^. Similar observations have been made in *E. coli*^53^ and *B. subtilis*^29^. This suggests that the *mnrS* riboswitch, when released from the full-length *mnrS-mntY* transcript, might act as a trans-acting sRNA. Further supporting this hypothesis is the presence of multiple 5’UTR-derived sRNAs across bacteria^54–56^. The first examples of riboswitches released by premature termination of transcription and shown to function as trans-acting RNA are the S-adenosylmethionine (SAM) riboswitches, SreA and SreB. Both modulate the synthesis of the master regulator of virulence PfrA in *Listeria monocytogenes*^57^. While the release and cellular accumulation of the *mnrS* riboswitch suggests secondary trans-acting functions, such roles have yet to be identified.

### Multiple functions contribute to the importance of MntY in S. aureus

The *mnrS* riboswitch and IsrR work together to regulate the synthesis of MntY, a Mn efflux pump which belongs to the TerC family (Figure 7C). The present study revealed that MntY protects against Mn intoxication, bringing the total number of known Mn efflux pumps in *S. aureus* to two. While MntY rescues a strain lacking MntE, the converse is not true, indicating that they possess non-redundant functions. Notably, the *mnrS-mntY* locus is expressed even in the absence of toxic concentrations of Mn (Figure 1B). While the synthesis of MntE is regulated by the transcriptional metal-dependent protein MntR^11^, the Mn-sensing riboswitch permits the synthesis of MntY at lower Mn concentration. The possibility that MntY contribute to physiology in the absence of toxicity is born out by the profound pleomorphic impacts that loss of MntY has on *S. aureus*. This includes a drastic reduction in the production of LTA (Figure 5D). MntY is a homolog of YkoY (MeeY) in *B. subtilis*, which was recently revealed to promote metalation of extracellular Mn-dependent proteins, including LTA synthase^36^. YkoY can complement the growth defect of a Δ*mnrS-mntY* mutant (Figure 5B). Thus, it seems likely that under non-toxic conditions, MntY contributes to metalation of extracellular Mn-dependent proteins in *S. aureus*. However, YkoY does not complement the ability of a Δ*mntE* mutant to grow in the presence of Mn toxic levels (Figure 5C), and YkoY appears to make only a minimal contribution to protect *B. subtilis* from Mn toxicity^35^. Taken together, these observations indicate that MntY and YkoY, while homologous, have non-overlapping roles. Indeed, MntY is involved in both the prevention of Mn toxicity and the metalation of essential proteins, whereas YkoY is primarily involved in the maintenance of metal-dependent processes (Figure 7A). *B. subtilis* appears to possess a significantly expanded repertoire of Mn efflux pumps when compared to *S. aureus*, having two Mn^2+^-inducible cation diffusion facilitator efflux pumps MneP and MneS^37^, an ABC-type exporter YknUV^58^ and two TerC family membrane protein paralogs, YceF and YkoY^35^. Notably, loss of both YkoY and YceF in *B. subtilis* is necessary to have the same effect on colony size and growth as loss of MntY in *S. aureus*^36^. YkoY homologs from *L. monocytogenes* and *B. anthracis* complement a *B. subtilis ykoY yceF* double mutant^36^, but it is unclear if they contribute to Mn detoxication. It seems likely that lifestyle and environment drive the differences in the efflux pump repertoire.

### MntY is essential for viability and virulence, but its expression must be tightly controlled

The contribution of MntY to maintaining the activity of LtaS, and consequently, the production of LTA, highlights the critical role of this protein in preserving critical processes. Hence, it is not surprising that loss of MntY has strong pleomorphic impacts including sensitizing *S. aureus* to antimicrobial compounds (calprotectin, daptomycin), immune cell killing and reducing the ability to causes infection (Figure 7B). At the same time, the activity of MntY must be tightly controlled as its overexpression also appears to be lethal. Disruption of cell wall synthesis, including via inactivation of LtaS, has been proposed as a promising avenue for therapeutic development given the extracellular nature of many targets and the critical contribution of the cell wall to stress resistance^59^. The current study suggests a new mechanism for disrupting cell wall synthesis, namely preventing enzymes such as LtaS from obtaining the cofactor critical for their activity, by inhibiting MntY. The intricate regulation of MntY to avoid the toxicity associated with its overexpression and its essential nature, suggest that it may have a reduced tolerance for mutation. This, combined with its surface localization, makes MntY an attractive target for therapeutic development and an opportunity to combat the redoubtable human pathogen *S. aureus*. TerC family proteins being conserved, this therapeutic strategy could be extended to other pathogens such as *L. monocytogenes* and *B. anthracis*^36^.

## MATERIAL AND METHODS

### Strains, plasmids, and growth conditions

Bacterial strains and plasmids used in this study are listed in Table S3A. For plasmids construction, PCR fragments amplified by oligonucleotides indicated in Table S3B were digested with respective restriction enzymes and ligated into a similarly digested vector. Plasmids isolated from *E. coli* TOP10 (Thermo Fischer Scientific) were then transferred into proper backgrounds. To efficiently transform *S. aureus* HG001 cells, *E. coli strain* IM08B^60^ was used as recipient strain for plasmid amplification. Unlike plasmid-based gene reporter systems used in *S. aureus* (pJB185), *lacZ* fusions were inserted as a single copy into the chromosome of *E. coli* MG1655 derivates^61^.

*E. coli* strains MG1655, TOP10 or IM08B and *B. subtilis* 168 were streaked onto LB agar plates and cultivated in lysogeny broth (LB) or M63 minimal medium. *S. aureus* strains were isolated on brain-heart infusion (BHI) or blood agar plates. Overnight cultures (3 mL) in BHI or in RPMI + 1% CA, a Roswell Park Memorial Institute (RPMI, Sigma-Aldrich) medium supplemented with 1% casamino acids (MP Biomedicals), were diluted 50-fold in fresh medium and grown with shaking at 37°C (180 rpm, 5:1 flask-to-medium ratio). To induce metal starvation, BHI or RPMI + 1% CA medium were mixed with 5% Chelex-100 resin (Sigma-Aldrich) during 6h at room temperature. The resulting media, BHI chelex and NRPMI, were then sterile filtered and, according to Kehl-Fie *et al.*^62^, were complemented with respective metal ions at the following concentrations: 1 mM MgCl_2_, 100 µM CaCl_2_, 25 µM ZnCl_2_, 25 µM MnCl_2_ and 1 µM FeSO_4_ (Sigma-Aldrich). The endogenous expression of *isrR* gene was induced by addition of the Fe chelator 2,2′-dipyridyl (DIP, 250 μM; Sigma-Aldrich). Antibiotics were added when required: 100 μg/mL of ampicillin (pJET, pMAD or pJB185 derivates in *E. coli*), 10 μg/mL of chloramphenicol (pJB185 derivates in *S. aureus*), 10 μg/mL of erythromycin (pMAD and pEW derivates in *S. aureus*), 300 µg/mL of rifampicin (half-life determination; *S. aureus*). Daptomycin hypersensitivity was tested in the presence of 0.5, 1 and 5 µg/mL of daptomycin (Sigma) and 100 µg/mL CaCl_2_.

Cells were streaked onto blood agar and 5% skim milk LB agar plates to determine haemolytic and proteolytic activities, respectively.

### Mutagenesis and complementation

The deletion of *isrR* gene in *S. aureus* HG001 strain was carried out using the pMAD vector^63^. Upstream and downstream regions of *isrR* gene (≈500 nts) were amplified by PCR (*isrR*-UF-XbaI/*isrR*-UR and *isrR*-DF/*isrR*-DR-XhoI, respectively) (Table S3B) and then cloned into XbaI/XhoI digested pMAD plasmid. The resulting plasmid was transferred to IM08B recipient strain and then electroporated into *S. aureus* HG001. As previously described^63^, growth at restrictive temperature (44°C) was followed by subcultures at 28°C to favor double crossover. The gene deletion was verified by PCR using oligonucleotides *isrR*-del-For and *isrR*-del-Rev. The same approach was used to delete *mntE, mnrS* and *mnrS*-*mntY* and to mutated *isrR* sequence (*isrR*-2G).

To complement mutant strains, *mntY*, *mnrS-mntY*, *mnrW-ykoY, mswM-yybP*, and *mntE* coding sequences were amplified using indicated primers (Table S3B) and cloned under the control of a constitutive BlaZ-derived promoter as described in Menendez-Gil *et al*^45^. All constructs were verified by DNA sequencing (Eurofins).

### In silico prediction of IsrR targets

A list of putative targets has been obtained using CopraRNA^64^ (default parameters) and the sequence of *isrR* genes from *Staphylococcus aureus* strain HG001 isolate RN1 (NZ_CP018205), *Staphylococcus epidermidis* RP62A (NC_002976), *Staphylococcus epidermidis* ATCC 12228 (NC_004461), *Staphylococcus haemolyticus* JCSC1435 (NC_007168), *Staphylococcus aureus* subsp. *aureus* NCTC 8325 (NC_007795), *Staphylococcus pseudintermedius* ED99 (NC_017568), *Staphylococcus warneri* SG1 (NC_020164.1), *Staphylococcus pasteuri* SP1 (NC_022737), *Staphylococcus capitis* subsp. *capitis* strain AYP1020 (NZ_CP007601), *Staphylococcus aureus* strain CA12 (NZ_CP007672), *Staphylococcus xylosus* strain SMQ-121 (NZ_CP008724), *Staphylococcus schleiferi* strain 1360-13 (NZ_CP009470), *Staphylococcus aureus* strain MS4 (NZ_CP009828), *Staphylococcus aureus* strain ZJ5499 (NZ_CP011685), Staphylococcus sp. AntiMn-1 (NZ_CP012968), *Staphylococcus equorum* strain KS1039 (NZ_CP013114), *Staphylococcus lugdunensis* strain K93G (NZ_CP017069), *Staphylococcus succinus* strain 14BME20 (NZ_CP018199), *Staphylococcus capitis* strain FDAARGOS_378 (NZ_CP023966), *Staphylococcus simiae* strain NCTC13838 (NZ_LT906460).

### Protein structure prediction and comparison

The sequence alignment was performed using Clustal Omega using HG001_00869 (MntY) from *S. aureus* HG001 and BSU_40560 (MeeY/YkoY) from *B. subtilis* 168. MntY secondary structure was predicted using AlphaFold^65^ and then aligned and superposed with MeeY structure (Uniprot accession O34997· YKOY_BACSU) from *B. subtillis* 168 using chimera software with default parameters^66^. The sequence alignment score is 841.8 and the root mean square deviation (RMSD) between 206 pruned atom pairs is 0.703 angstroms (across all 247 pairs: 4.318).

### Gel retardation assays

Transcription start sites were determined according to Koch et al.^67^. PCR fragments containing T7-*mntE* (full-length, from −44 to +867), T7-*yybP-ykoY-mntP* (from −225 to +50, *E. coli*) and T7-*sraF-alx* (from −206 to +170, *E. coli*) were used for as DNA template for in vitro transcription with T7 RNA polymerase. T7-*isrR* (full-length), T7-*mnrS* (full-length), T7-*mnrS-mntY* (from −183 to +115), T7-*ryhB* (full-length, *E. coli*), T7-*fsrA* (full-length, *B. subtilis*), T7-*mswM-yybP* (from −307 to +172, *B. subtilis*), and T7-*mnrW-ykoY* (from −21 to +289, *B. subtilis*) were cloned into pJET1.2*/*blunt plasmid (ThermoFisher). The *isrG*-2G and *mnrS*-2C mutants were obtained using the QuickChange site-directed mutagenesis method (Agilent Technologies) following manufacturer’s instructions. Oligonucleotides containing the desired mutations are listed in Table S3B. pJET-T7-*isrR* and pJET-T7-*mnrS* were used as template to generate the mutated plasmids. T7 transcriptions were performed using XbaI and/or XhoI-digested plasmids as template. RNAs were finally purified and radiolabeled when required^26^. 5’-radiolabelled IsrR, IsrR-2G, *mnrS*, FsrA or RyhB (20,000 cpm/sample, concentration <1 pM) and above-mentioned cold RNAs were separately denatured at 90°C in the buffer GR- (20 mM Tris-HCl pH 7.5, 60 mM KCl, 40 mM NH_4_Cl, 3 mM DTT), cooled 1 min on ice, and incubated at room temperature for 15 min in presence of 10 mM MgCl_2_. Renatured RNAs were then mix and incubated at 37°C for 15 min. Finally, samples were loaded on a 6% polyacrylamide gel under non-denaturing conditions (300 V, 4°C). ImageJ software^68^ was used to perform densitometric analysis. Results are representative of two independent experiments.

### Probing experiments

PbAc and RNase T1 probing assays were performed as previously described^69^. Briefly, 0.04 µM of 5’-end-labeled *mnrS* (full-length) transcript was incubated with (+) or without (-) 1 µM of IsrR/IsrR-2G sRNA. Radiolabeled RNA was incubated at 90°C for 5 min with alkaline hydrolysis buffer (OH; Ambion) or at 37°C for 5 min with ribonuclease T1 (0.1 U; Ambion) to obtain the OH ladder and the T1 ladder, respectively.

### Termination efficiency assays

Termination efficiency assays were performed as described in Salvail et al.^70^ with slight modifications. 100 nM of linear DNA template (P*lysC*-*mnrS*; Table S3B) were mixed with increasing concentration of MnCl_2_ (0 to 10 mM; final volume of 6 µL). 3 µL of transcription master mix, which is composed of 0.9 µL of 10X transcription buffer (200 mM Tris-Cl pH 8.0, 1 M KCl, 20 mM MgCl_2_, 1 mM EDTA), 0.9 µL of 10X GTP-free initiation mix (25 µM CTP, 25 µM ATP and 10 µM UTP), 0.1 µL of 1 mg/mL BSA, 0.13 µL of 10 mM ApA (Jena Bioscience), 0.2 µL of 50% glycerol, 2 µCi α-UTP, and 0.4 U of *E. coli* RNA polymerase (Holoenzyme, NEB) were added. The initiation of transcription was performed at 37°C during 10 min. Here, the RNA polymerase stalled at the first G residue. Then, 1 µL of 10X elongation mix (1.5 mM GTP, 1.5 mM ATP, 1.5 mM CTP, 0.1 mM UTP, 1 mg/mL heparin in 1X transcription buffer) was added to resume the reaction. Samples were incubated 30 min at 37°C. Reactions were finally stopped by addition of 10 µL of loading buffer II (95% Formamide, 18 mM EDTA, 0.025% SDS, 0.025% xylene cyanol, and 0.025% bromophenol blue), and resolved on 10% polyacrylamide gel.

### Growth monitoring assays

Freshly streaked bacterial colonies were used to inoculate overnight cultures incubated at 37°C with constant shaking (180 rpm). Cultures were diluted (OD_600nm_=0.05) in fresh media supplemented with 0, 40, and 400 µM MnCl_2_ before being placed in a 96-well flat-bottomed plate and then incubated at 37°C with constant agitation for 10-20 hours. Growth was measured at OD_600nm_ in a microplate reader Multiskan SkyHigh multiplate spectrophotometer (ThermoFisher). At least three independent experiments were performed.

### Northern blot analysis

10 mL of bacterial suspension were collected at indicated OD_600nm_ or time. Bacterial pellets were resuspended in the RNA Pro Solution provided in the FastRNA Pro Blue kit (MP Biomedicals). The FastPrep apparatus (MP Biomedicals) was used to mechanically lyse the cells. Total RNA extraction was performed following manufacturer’s instructions.

10-20 µg total RNA were loaded on a 1% agarose gel containing 25 mM guanidium thiocyanate (Sigma-Aldrich) and electrophoresed at 120 V. RNAs were then transferred on Hybond N+ nitrocellulose membrane (GE Healthcare Life Sciences). Membranes were hybridized with specific digoxygenin (DIG)-labelled probes targeting RNA of interest or 5S rRNA (loading control), obtained using the DIG RNA Labelling kit (Roche) and corresponding oligonucleotides (Table S3B). The luminescent detection was carried out as previously described using CDP-Star (Roche)^71^. Results are representative of at least two independent experiments.

For the determination of RNA half-lives, 250 µM of DIP were added for 15 minutes to induce *isrR* expression. 10 mL of culture were collected before (0) and after (0.5, 1, 2.5, 5, 7.5, 10, 15, and 30 minutes) rifampicin treatment (300 µg/mL). Densitometric analyses were performed using Image J software^68^. Each experiment was performed at least twice.

### β-galactosidase assays

Constructs were obtained by PCR and cloned into pJB185 plasmid^72^. The MntP+45-LacZ and Alx+15-LacZ translational fusions were constructed using λ red-mediated recombineering. Fragments extended from −225 to +45 (*mntP*) or −206 to +15 (*alx*) relative to the ATG were inserted into NM580 strain, which contains pBAD-*ccdB* upstream of *lacZ* at the chromosomal *lac* locus^73^. These constructions were moved into new backgrounds by P1 transduction.

Bacterial cells from an overnight culture were diluted in 50 mL (flask; *E. coli*) or 10 mL of fresh medium (50 mL tubes; *S. aureus*). Cells were grown at 37°C and harvested at the appropriate OD_600nm_. When required, the production of sRNAs was induced by addition of 0.1% arabinose (*E. coli*). β-galactosidase kinetic measurements were performed at OD_420nm_ on a Multiskan SkyHigh multiplate spectrophotometer (ThermoFisher) and analyzed with SkanIt™ Software for Microplate Readers (version 6.1). The β-galactosidase activity was determined by applying the formula Vmax/OD_600nm_^74^. Processed data correspond to the mean of at least three independent experiments ± standard deviation (SD).

### LTA detection by Western blot

Lipoteichoic acid (LTA) detection was performed as previously described by He et al.^36^ with some changes. Briefly, cultures were grown at 37°C in agitation until the exponential phase (OD_600nm_=0.6). 5 mL of cultures were pelleted and washed with buffer I (100 mM Tris-HCl pH 7.4 with Halt™ protease inhibitor cocktail (1X); Thermo Scientific™). Cell pellets were resuspended in one volume of buffer I and one volume of 2X Laemmli buffer (without reducing agent and bromophenol Blue). Samples were then heated at 100 °C for 45LJmin, cooled down on ice for 5 min, and treated with 10 U/mL DNase I (Roche) at 37 °C for 30LJmin. After 5 min of centrifugation, supernatants were quantified with Pierce BCA protein assay (Thermo Scientific™) and stored at −20 °C for posterior analysis. β-mercaptoethanol and bromophenol blue were added before denaturation at 95 °C for 10 min. Samples were subsequently loaded (5 µg) onto a 20% SDS-PAGE polyacrylamide gel for electrophoretic separation. After transfer onto a PVDF membrane using a Trans-Blot Turbo Transfer System (Bio-Rad), the membrane was blocked with a PBS solution containing 3% BSA (w/v) for 30 min with shaking. The LTA was detected by using a Gram-positive LTA monoclonal antibody at 1:1000 dilution (Thermo Fisher, overnight incubation at 4°C) and a horseradish peroxidase (HRP)-conjugated Goat anti-mouse antibody 1:10,000 dilutions (BioRad, 1h incubation 4°C). After three washes in PBS 0.1% tween, the membrane was incubated with ECL reagents (Bio-Rad) and scanned with a ChemiDoc image system (Bio-Rad). As a loading control, we loaded another gel for Coomassie blue staining, and the same transferred membrane was stripped and re-probed for *Staphylococcus* Protein A detection with HRP-conjugated Anti-rabbit (Bio-Rad) at 1/3000 dilution.

### RNA-RNA interaction prediction

#### Riboswitch homologs, sRNA homologs and riboswitch clustering

The *yybP-ykoY* riboswitch dataset contains the respective Rfam sequences (RF00080)^75^, the sequences described in Dambach et al.^29^ and sequences which were additionally detected by a GLASSgo^76^ search with the *S. aureus* (CP125863.1:997786-997951) and *Bacillus* (CP000813.4:1297230-1297337) riboswitch sequences as queries. GLASSgo was done on bacterial genomes (https://zenodo.org/record/1320180) (E-value≤5, minimum sequence identity 57%, londen mode off). Sequences with overlapping genomic coordinates were merged and evaluated with Infernals cmsearch^77^ against the RF00080 model and a custom covariance model for riboswitches in front of the *alx* gene, using default parameters (E-value <= 0.01), resulting in 8,644 homolog sequences. The same procedure was applied to detect homologs of the Fe stress related sRNAs ArrF^41^ (2788, RF00444), FsrA^21^ (726, RF02273), Isar1^42^ (57, no Rfam family), IsrR^22^ (1905, RF02578), NrrF^43^ (515, RF01416), PrrF1/PrrF2^20^ (2695 & 2797, RF00444) and RyhB^19^ (3416, RF00057). ArrF, PrrF1 and PrrF2 share the same Rfam family and detected sequences might be partly redundant. Riboswitch homologs were clustered based on the annotated gene 3-prime of the riboswitch location. The respective amino acid sequences were clustered with mmseq2^78^ (min-seq-id = 0.2, C = 0.8, cov-mode = 1, cluster-mode = 2), initially resulting in 132 clusters, including 92 singleton clusters. The singleton and low member size cluster are partly likely due to a still incomplete set of existing riboswitch homologs as well es clustering and annotation artifacts. The initial clusters were manually curated to account for inconsistent genome annotations and a too sensitive clustering. Singleton clusters were not further investigated, which finally resulted in 16 riboswitch families.

#### Interaction predictions

Riboswitch homologs were aligned with Clustal Omega^79^ and the aligned sequences were trimmed to a common 5’end prior to interaction prediction. The prediction of potential interactions between all riboswitch homologs, sRNAs and sRNA homologs within a respective organism was done with IntaRNA^44^ (default parameters beside seedBP = 5). Based on the experimental, results we categorized the detected interaction sites based on their position in the riboswitch, i.e., interactions at the 5’ end that include the “GGGGAGUA” (5’GGGG_sites) motif and predicted interactions sites at other positions. All 5’GGGG_sites with a predicted interaction energy≤-9 kcal/mol were considered as potential positive sites. A frequently predicted interaction site between the 3’ end of the riboswitch and RyhB sRNA was identified in many Enterobacteria (Figure 4C). This pairing site was not considered, as no shift between RyhB and both riboswitches was observed in *E. coli* (Figure S4C).

#### Tree generation

The sRNA and riboswitch distribution are plotted across a Neighbor-Joining tree (Biopython - Bio.phylo) based on a Clustal Omega^79^ 16S rDNA alignment. An archeal 16S rDNA sequence from *Desulfurococcus* DSM2162 was used as outgroup for tree rooting. The trees are visualized with iTOL^80^. The outer ring shows the occurrence patterns of the 9 most frequent riboswitch families as bars. A gradient from dark colors to light colors (predicted interaction energy: ≥-22.19 kcal/mol to ≤-9 kcal/mol) indicates the predicted interaction potential of the respective riboswitch with one of the present sRNAs. If an organism harbors more than one sRNA, the lowest energy interaction is displayed. The gray color indicates an interaction energy >9 kcal/mol or no predicted 5’GGGG_site (definition see above). The inner ring shows the detected sRNAs. The rDNA sequences were extracted from the GenBank files with barrnap (https://github.com/tseemann/barrnap). For the remaining group, closely related organisms were condensed to one representative based on the 16S rDNA similarity (distance threshold = 10^-5^). Clades with very long branches were collapsed at an internal node (marked in red).

### Fluorescein-isothiocyanate (FITC) labeling

Bacteria from fresh cultures were prewashed in 0.1 M sodium bicarbonate buffer, pH 9.0. *S. aureus* cells were pelleted, and stained with 0,5µg/µL of fluorescein isothiocyanate (FITC; Invitrogen). Bacteria were incubated for 20 min at room temperature and washed three times in DPBS to remove unbound dye. Image acquisition was performed on a fluorescent microscope at 40 X (Leica™ TCS SP5). Images were processed using Image J software^68^.

### Calprotectin growth assays

Calprotectin (CP) growth assays were performed as described previously^81^. Briefly, overnight cultures (grown 16-18 h in BHI medium at 37°C) were diluted 1:100 into 96-well round-bottom plates containing 100 µL of growth medium (38% 3x BHI and 62% calprotectin buffer (20 mM Tris pH 7.5, 100 mM NaCl, 3 mM CaCl_2_)) in presence of varying concentrations of CP. For all assays, the bacteria were incubated with orbital shaking (180 rpm) at 37°C and growth was measured by assessing optical density (OD_600nm_) every 2 hours. Prior to measuring optical density, the 96-well plates were vortexed.

### Neutrophil survival assay

All participants provided written, informed consent in accordance with the principles outlined in the Declaration of Helsinki. Peripheral human blood samples were obtained from healthy donors through the Etablissement Français du Sang (EFS) in Strasbourg, France under authorization number ALC/PIL/DIR/AJR/FO/606. Samples were collected in tubes containing 3.8% sodium citrate. Primary human polymorphonuclear leukocytes (PMNs) were isolated using Percoll gradient separation as previously described^82^. Freshly prepared neutrophils were resuspended at a concentration of 1 × 10^7^ cells/mL and prewarmed at 37°C for 10 minutes. S. aureus cells were grown in BHI medium and harvested at an OD_600nm_ ≈ 1. A multiplicity of infection (MOI) of 5 (5:1) was used. Samples were incubated at 37°C with gentle rotation (80 rpm) for 20 min. (A) Bacterial survival was assessed by plating serial 10-fold dilutions on BHI agar plates and quantified as colony-forming units per milliliter (CFU/mL) (n=8; 3 blood donors). Alternatively, (B) samples were centrifugated at 300 g to separate internalized from extracellular bacteria (supernatant). Pellets were washed with DPBS and then incubated for 20 minutes in RPMI media containing 25 µg/mL lysostaphin to eliminate remaining extracellular bacteria. After centrifugation, cell pellets were resuspended in sterile Triton 0.01% and allowed to stand at room temperature for 5-10 min. Extracellular and internalized bacterial loads were then assessed by plating serial 10-fold dilutions on BHI agar plates (n=4; 1 blood donor).

### THP-1 derived macrophage infection

THP-1 cells were resuspended in RPMI 1640 supplemented with 2.5% fetal bovine serum (FBS) and plated 48 h before infection at a density of 5LJ×LJ10^5^ cells per well. THP-1 cells were differentiated into macrophage-like cells by treatment with 100 ng/ml phorbol 12-myristate 13-acetate PMA (Sigma-Aldrich). Overnight bacterial growths were diluted 1:100 in BHI medium and grown until an OD_600nm_ ≈ 1. Infections were performed at a multiplicity of infection (MOI) of 5 (5:1). Cells were centrifuged at 300 g, at room temperature for 4LJmin, and incubated at 37 °C in a 5% CO_2_ humidified atmosphere for 90 min. (A) Bacterial survival was assessed by plating serial 10-fold dilutions on BHI agar plates and quantified as CFU/mL (n=7). Alternatively, (B) samples were centrifugated at 300 g to separate internalized from extracellular bacteria (supernatant). After centrifugation, remaining extracellular bacteria were lysed by addition of 25 µg/ml lysostaphin and 150 µg/ml gentamicin (Sigma-Aldrich) for 1h. As previously mentioned, extracellular and internalized bacterial loads were assessed by plating serial 10-fold dilutions on BHI agar plates (n=7).

### *S. aureus* mouse infection

Animal protocols were approved by the animal welfare ethics committee “C2EA-35” (Protocol APAFIS #44185-2023111310376755). Bacterial cultures were started from freshly isolated single colonies and grown overnight in BHI (3 mL in 15 mL sterile tubes) at 37°C, shaking at 180 rpm. On the next day, a 1:100 subculture in 100 mL fresh BHI (in 500 mL sterile flasks) was incubated at 37°C with shaking at 180 rpm until OD_600nm_ ≈ 1 for the WT and 0.6 for the Δ*mnrS-mntY* mutant. Cultures were then centrifuged, and cell pellets were washed twice in cold DPBS. Cells were finally resuspended in sterile DPBS to obtain an OD_600nm_ between 1.15-1.25 for the WT and 2.20-2.50 for the Δ*mnrS-mntY* mutant, corresponding to 1×10^8^ CFU per mL. Two groups of 6 mice (female 9-to-10-week-old C57BL/6J mice; Charles River Laboratories) were inoculated with 1 × 10^7^ CFU of bacteria in 100 µL by retroorbital injection. Mice were weighed and monitored daily. At 24 and 72h post-infection, mice were euthanized, and organs such as kidney, spleen, liver, heart, and liver were harvested and bacterial burdens determined by plating on BHI agar to determine CFU/organ. Colonies were enumerated after incubation at 37 °C for 24LJh.

## ACKNOWLEDGMENTS

The authors would like to thank Dr. Jeffrey Bose for its generous gift of the pJB185 vector. They would also like to thank Dr. Kevin Waldron and all team members for helpful discussions. D.L. was supported by the Agence Nationale de la Recherche (ANR; grant ANR-20-CE12-0021, MetalAureus). J.G. was supported by the Deutsche Forschungsgemeinschaft (grant DFG GE 3159/1-1). D.L. and T.K.F. were supported by Thomas Jefferson Fund, a program of FACE Foundation launched in collaboration with the French Embassy. The work of the Interdisciplinary Thematic Institute IMCBio, as part of the ITI 2021-2028 program of the University of Strasbourg, CNRS and INSERM, was supported by IdEx Unistra (ANR-10-IDEX-0002), by SFRI-STRAT’US project (ANR 20-SFRI-0012), and by EUR IMCBio (IMCBio ANR-17-EURE-0023) under the framework of the French Investments for the Future Program. Work in the laboratory of T.K.F. is supported by grants from the NIH (R01AI179695) and by the University of Iowa’s Year 2 P3 Strategic Initiatives Program through funding received for the project entitled “High Impact Hiring Initiative (HIHI): A Program to Strategically Recruit and Retain Talented Faculty.”

## AUTHOR CONTRIBUTIONS

Conceptualization, G.G.E. and D.L.; Methodology, G.G.E, T.K.F., J.G., D.L.; Software, F.H., and J.G.; Validation, G.G.E., K.P., J.G., and D.L.; Investigation, G.G.E., K.P., F.H., J.N.R., C.D.V.V, J.M, M.V.B, J.G. and D.L.; Writing – Original Draft, G.G.E. and D.L.; Writing – Review & Editing, G.G.E., K.P., B.M., E.M., P.R., T.K.F, J.G., and D.L.; Visualization, G.G.E., K.P., F.H., J.N.R., J.G., and D.L.; Supervision, E.M., T.K.F., J.G., and D.L.; Project Administration, D.L.; Funding Acquisition, P.R., T.K.F, J.G., and D.L.

## REFERENCES

1. Ullah, I., and Lang, M. (2023). Key players in the regulation of iron homeostasis at the host-pathogen interface. Frontiers in Immunology 14. 10.3389/fimmu.2023.1279826.

2. Murdoch, C.C., and Skaar, E.P. (2022). Nutritional immunity: the battle for nutrient metals at the host-pathogen interface. Nat Rev Microbiol 20, 657–670. 10.1038/s41579-022-00745-6.

3. Kelliher, J.L., and Kehl-Fie, T.E. (2016). Competition for Manganese at the Host-Pathogen Interface. Progress in molecular biology and translational science 142, 1–25. 10.1016/bs.pmbts.2016.05.002.

4. Sheldon, J.R., and Skaar, E.P. (2019). Metals as phagocyte antimicrobial effectors. Current opinion in immunology 60, 1–9. 10.1016/j.coi.2019.04.002.

5. Djoko, K.Y., Ong, C.L., Walker, M.J., and McEwan, A.G. (2015). The Role of Copper and Zinc Toxicity in Innate Immune Defense against Bacterial Pathogens. J Biol Chem 290, 18954–18961. 10.1074/jbc.R115.647099.

6. O’Neal, S.L., and Zheng, W. (2015). Manganese Toxicity Upon Overexposure: a Decade in Review. Current environmental health reports 2, 315–328. 10.1007/s40572-015-0056-x.

7. Schroeder, H.A., Balassa, J.J., and Tipton, I.H. (1966). Essential trace metals in man: manganese. A study in homeostasis. Journal of chronic diseases 19, 545–571. 10.1016/0021-9681(66)90094-4.

8. Jordan, M.R., Wang, J., Capdevila, D.A., and Giedroc, D.P. (2020). Multi-metal nutrient restriction and crosstalk in metallostasis systems in microbial pathogens. Curr Opin Microbiol 55, 17–25. 10.1016/j.mib.2020.01.010.

9. Hossain, S., Morey, J.R., Neville, S.L., Ganio, K., Radin, J.N., Norambuena, J., Boyd, J.M., McDevitt, C.A., and Kehl-Fie, T.E. (2023). Host subversion of bacterial metallophore usage drives copper intoxication. mBio 14, e0135023. 10.1128/mbio.01350-23.

10. Al-Tameemi, H., Beavers, W.N., Norambuena, J., Skaar, E.P., and Boyd, J.M. (2021). Staphylococcus aureus lacking a functional MntABC manganese import system has increased resistance to copper. Mol Microbiol 115, 554–573. 10.1111/mmi.14623.

11. Grunenwald, C.M., Choby, J.E., Juttukonda, L.J., Beavers, W.N., Weiss, A., Torres, V.J., and Skaar, E.P. (2019). Manganese Detoxification by MntE Is Critical for Resistance to Oxidative Stress and Virulence of Staphylococcus aureus. mBio 10. 10.1128/mBio.02915-18.

12. Horsburgh, M.J., Wharton, S.J., Cox, A.G., Ingham, E., Peacock, S., and Foster, S.J. (2002). MntR modulates expression of the PerR regulon and superoxide resistance in Staphylococcus aureus through control of manganese uptake. Mol Microbiol 44, 1269–1286. 10.1046/j.1365-2958.2002.02944.x.

13. Troxell, B., and Hassan, H.M. (2013). Transcriptional regulation by Ferric Uptake Regulator (Fur) in pathogenic bacteria. Front Cell Infect Microbiol 3, 59. 10.3389/fcimb.2013.00059.

14. Chandrangsu, P., Rensing, C., and Helmann, J.D. (2017). Metal homeostasis and resistance in bacteria. Nat Rev Microbiol 15, 338–350. 10.1038/nrmicro.2017.15.

15. Charbonnier, M., González-Espinoza, G., Kehl-Fie, T.E., and Lalaouna, D. (2022). Battle for Metals: Regulatory RNAs at the Front Line. Front Cell Infect Microbiol 12, 952948. 10.3389/fcimb.2022.952948.

16. Chareyre, S., and Mandin, P. (2018). Bacterial Iron Homeostasis Regulation by sRNAs. Microbiology spectrum 6. 10.1128/microbiolspec.RWR-0010-2017.

17. Papenfort, K., and Melamed, S. (2023). Small RNAs, Large Networks: Posttranscriptional Regulons in Gram-Negative Bacteria. Annual review of microbiology 77, 23–43. 10.1146/annurev-micro-041320-025836.

18. Desgranges, E., Marzi, S., Moreau, K., Romby, P., and Caldelari, I. (2019). Noncoding RNA. Microbiology spectrum 7. 10.1128/microbiolspec.GPP3-0038-2018.

19. Masse, E., and Gottesman, S. (2002). A small RNA regulates the expression of genes involved in iron metabolism in Escherichia coli. Proc Natl Acad Sci U S A 99, 4620–4625. 10.1073/pnas.032066599.

20. Wilderman, P.J., Sowa, N.A., FitzGerald, D.J., FitzGerald, P.C., Gottesman, S., Ochsner, U.A., and Vasil, M.L. (2004). Identification of tandem duplicate regulatory small RNAs in Pseudomonas aeruginosa involved in iron homeostasis. Proc Natl Acad Sci U S A 101, 9792–9797. 10.1073/pnas.0403423101.

21. Gaballa, A., Antelmann, H., Aguilar, C., Khakh, S.K., Song, K.B., Smaldone, G.T., and Helmann, J.D. (2008). The Bacillus subtilis iron-sparing response is mediated by a Fur-regulated small RNA and three small, basic proteins. Proc Natl Acad Sci U S A 105, 11927–11932. 10.1073/pnas.0711752105.

22. Coronel-Tellez, R.H., Pospiech, M., Barrault, M., Liu, W., Bordeau, V., Vasnier, C., Felden, B., Sargueil, B., and Bouloc, P. (2022). sRNA-controlled iron sparing response in Staphylococci. Nucleic Acids Res 50, 8529–8546. 10.1093/nar/gkac648.

23. Barrault, M., Chabelskaya, S., Coronel-Tellez, R.H., Toffano-Nioche, C., Jacquet, E., and Bouloc, P. (2024). Staphylococcal aconitase expression during iron deficiency is controlled by an sRNA-driven feedforward loop and moonlighting activity. Nucleic Acids Res. 10.1093/nar/gkae506.

24. Rios-Delgado, G., McReynolds, A.K.G., Pagella, E.A., Norambuena, J., Briaud, P., Zheng, V., Munneke, M.J., Kim, J., Racine, H., Carroll, R., et al. (2024). The Staphylococcus aureus small non-coding RNA IsrR regulates TCA cycle activity and virulence. bioRxiv.

25. Ganske, A., Busch, L.M., Hentschker, C., Reder, A., Michalik, S., Surmann, K., Völker, U., and Mäder, U. (2024). Exploring the targetome of IsrR, an iron-regulated sRNA controlling the synthesis of iron-containing proteins in Staphylococcus aureus. Front Microbiol 15, 1439352. 10.3389/fmicb.2024.1439352.

26. Lalaouna, D., Baude, J., Wu, Z., Tomasini, A., Chicher, J., Marzi, S., Vandenesch, F., Romby, P., Caldelari, I., and Moreau, K. (2019). RsaC sRNA modulates the oxidative stress response of Staphylococcus aureus during manganese starvation. Nucleic Acids Research 47, 9871–9887. 10.1093/nar/gkz728.

27. Garcia, Y.M., Barwinska-Sendra, A., Tarrant, E., Skaar, E.P., Waldron, K.J., and Kehl-Fie, T.E. (2017). A Superoxide Dismutase Capable of Functioning with Iron or Manganese Promotes the Resistance of Staphylococcus aureus to Calprotectin and Nutritional Immunity. PLoS Pathog 13, e1006125. 10.1371/journal.ppat.1006125.

28. Kavita, K., and Breaker, R.R. (2023). Discovering riboswitches: the past and the future. Trends in biochemical sciences 48, 119–141. 10.1016/j.tibs.2022.08.009.

29. Dambach, M., Sandoval, M., Updegrove, Taylor B., Anantharaman, V., Aravind, L., Waters, Lauren S., and Storz, G. (2015). The Ubiquitous yybP-ykoY Riboswitch Is a Manganese-Responsive Regulatory Element. Molecular cell 57, 1099–1109.

30. Price, Ian R., Gaballa, A., Ding, F., Helmann, John D., and Ke, A. (2015). Mn2+-Sensing Mechanisms of yybP-ykoY Orphan Riboswitches. Molecular cell 57, 1110–1123. 10.1016/j.molcel.2015.02.016.

31. Barrick, J.E., Corbino, K.A., Winkler, W.C., Nahvi, A., Mandal, M., Collins, J., Lee, M., Roth, A., Sudarsan, N., Jona, I., et al. (2004). New RNA motifs suggest an expanded scope for riboswitches in bacterial genetic control. Proc Natl Acad Sci U S A 101, 6421–6426. 10.1073/pnas.0308014101.

32. Zeinert, R., Martinez, E., Schmitz, J., Senn, K., Usman, B., Anantharaman, V., Aravind, L., and Waters, L.S. (2018). Structure-function analysis of manganese exporter proteins across bacteria. The Journal of biological chemistry 293, 5715–5730. 10.1074/jbc.M117.790717.

33. Suddala, K.C., Price, I.R., Dandpat, S.S., Janecek, M., Kuhrova, P., Sponer, J., Banas, P., Ke, A., and Walter, N.G. (2019). Local-to-global signal transduction at the core of a Mn(2+) sensing riboswitch. Nature communications 10, 4304. 10.1038/s41467-019-12230-5.

34. Waters, L.S., Sandoval, M., and Storz, G. (2011). The Escherichia coli MntR miniregulon includes genes encoding a small protein and an efflux pump required for manganese homeostasis. J Bacteriol 193, 5887–5897. 10.1128/jb.05872-11.

35. Paruthiyil, S., Pinochet-Barros, A., Huang, X., and Helmann, J.D. (2020). Bacillus subtilis TerC Family Proteins Help Prevent Manganese Intoxication. Journal of bacteriology 202, e00624–00619. 10.1128/JB.00624-19.

36. He, B., Sachla, A.J., and Helmann, J.D. (2023). TerC proteins function during protein secretion to metalate exoenzymes. Nature communications 14, 6186. 10.1038/s41467-023-41896-1.

37. Huang, X., Shin, J.H., Pinochet-Barros, A., Su, T.T., and Helmann, J.D. (2017). Bacillus subtilis MntR coordinates the transcriptional regulation of manganese uptake and efflux systems. Mol Microbiol 103, 253–268. 10.1111/mmi.13554.

38. Mäder, U., Nicolas, P., Depke, M., Pané-Farré, J., Debarbouille, M., van der Kooi-Pol, M.M., Guérin, C., Dérozier, S., Hiron, A., Jarmer, H., et al. (2016). Staphylococcus aureus Transcriptome Architecture: From Laboratory to Infection-Mimicking Conditions. PLoS Genet 12, e1005962. 10.1371/journal.pgen.1005962.

39. Jumper, J., Evans, R., Pritzel, A., Green, T., Figurnov, M., Ronneberger, O., Tunyasuvunakool, K., Bates, R., Žídek, A., Potapenko, A., et al. (2021). Highly accurate protein structure prediction with AlphaFold. Nature 596, 583–589. 10.1038/s41586-021-03819-2.

40. Bastet, L., Bustos-Sanmamed, P., Catalan-Moreno, A., Caballero, C.J., Cuesta, S., Matilla-Cuenca, L., Villanueva, M., Valle, J., Lasa, I., and Toledo-Arana, A. (2022). Regulation of Heterogenous LexA Expression in Staphylococcus aureus by an Antisense RNA Originating from Transcriptional Read-Through upon Natural Mispairings in the sbrB Intrinsic Terminator. Int J Mol Sci 23. 10.3390/ijms23010576.

41. Jung, Y.S., and Kwon, Y.M. (2008). Small RNA ArrF regulates the expression of sodB and feSII genes in Azotobacter vinelandii. Current microbiology 57, 593–597. 10.1007/s00284-008-9248-z.

42. Georg, J., Kostova, G., Vuorijoki, L., Schön, V., Kadowaki, T., Huokko, T., Baumgartner, D., Müller, M., Klähn, S., Allahverdiyeva, Y., et al. (2017). Acclimation of Oxygenic Photosynthesis to Iron Starvation Is Controlled by the sRNA IsaR1. Curr Biol 27, 1425–1436.e1427. 10.1016/j.cub.2017.04.010.

43. Mellin, J.R., Goswami, S., Grogan, S., Tjaden, B., and Genco, C.A. (2007). A novel fur- and iron-regulated small RNA, NrrF, is required for indirect fur-mediated regulation of the sdhA and sdhC genes in Neisseria meningitidis. J Bacteriol 189, 3686–3694. 10.1128/jb.01890-06.

44. Mann, M., Wright, P.R., and Backofen, R. (2017). IntaRNA 2.0: enhanced and customizable prediction of RNA-RNA interactions. Nucleic Acids Res 45, W435–w439. 10.1093/nar/gkx279.

45. Menendez-Gil, P., Caballero, C.J., Catalan-Moreno, A., Irurzun, N., Barrio-Hernandez, I., Caldelari, I., and Toledo-Arana, A. (2020). Differential evolution in 3’UTRs leads to specific gene expression in Staphylococcus. Nucleic Acids Res 48, 2544–2563. 10.1093/nar/gkaa047.

46. Bronesky, D., Desgranges, E., Corvaglia, A., Francois, P., Caballero, C.J., Prado, L., Toledo-Arana, A., Lasa, I., Moreau, K., Vandenesch, F., et al. (2019). A multifaceted small RNA modulates gene expression upon glucose limitation in Staphylococcus aureus. The EMBO journal 38. 10.15252/embj.201899363.

47. Wörmann, M.E., Corrigan, R.M., Simpson, P.J., Matthews, S.J., and Gründling, A. (2011). Enzymatic activities and functional interdependencies of Bacillus subtilis lipoteichoic acid synthesis enzymes. Mol Microbiol 79, 566–583. 10.1111/j.1365-2958.2010.07472.x.

48. Bastet, L., Korepanov, A.P., Jagodnik, J., Grondin, J.P., Lamontagne, A.M., Guillier, M., and Lafontaine, D.A. (2024). Riboswitch and small RNAs modulate btuB translation initiation in Escherichia coli and trigger distinct mRNA regulatory mechanisms. Nucleic Acids Res 52, 5852–5865. 10.1093/nar/gkae347.

49. Desgranges, E., Barrientos, L., Herrgott, L., Marzi, S., Toledo-Arana, A., Moreau, K., Vandenesch, F., Romby, P., and Caldelari, I. (2022). The 3’UTR-derived sRNA RsaG coordinates redox homeostasis and metabolism adaptation in response to glucose-6-phosphate uptake in Staphylococcus aureus. Mol Microbiol 117, 193–214. 10.1111/mmi.14845.

50. Melamed, S., Peer, A., Faigenbaum-Romm, R., Gatt, Y.E., Reiss, N., Bar, A., Altuvia, Y., Argaman, L., and Margalit, H. (2016). Global Mapping of Small RNA-Target Interactions in Bacteria. Molecular cell 63, 884–897. 10.1016/j.molcel.2016.07.026.

51. Rodgers, M.L., O’Brien, B., and Woodson, S.A. (2023). Small RNAs and Hfq capture unfolded RNA target sites during transcription. Mol Cell 83, 1489–1501.e1485. 10.1016/j.molcel.2023.04.003.

52. Roberts, C., Anderson, K.L., Murphy, E., Projan, S.J., Mounts, W., Hurlburt, B., Smeltzer, M., Overbeek, R., Disz, T., and Dunman, P.M. (2006). Characterizing the effect of the Staphylococcus aureus virulence factor regulator, SarA, on log-phase mRNA half-lives. J Bacteriol 188, 2593–2603. 10.1128/jb.188.7.2593-2603.2006.

53. Argaman, L., Hershberg, R., Vogel, J., Bejerano, G., Wagner, E.G., Margalit, H., and Altuvia, S. (2001). Novel small RNA-encoding genes in the intergenic regions of Escherichia coli. Curr Biol 11, 941–950.

54. Adams, P.P., Baniulyte, G., Esnault, C., Chegireddy, K., Singh, N., Monge, M., Dale, R.K., Storz, G., and Wade, J.T. (2021). Regulatory roles of Escherichia coli 5’ UTR and ORF-internal RNAs detected by 3’ end mapping. eLife 10, e62438. 10.7554/eLife.62438.

55. Cengher, L., Manna, A.C., Cho, J., Theprungsirikul, J., Sessions, K., Rigby, W., and Cheung, A.L. (2022). Regulation of neutrophil myeloperoxidase inhibitor SPIN by the small RNA Teg49 in Staphylococcus aureus. Mol Microbiol 117, 1447–1463. 10.1111/mmi.14919.

56. Thomason, M.K., Voichek, M., Dar, D., Addis, V., Fitzgerald, D., Gottesman, S., Sorek, R., and Greenberg, E.P. (2019). A rhlI 5’ UTR-Derived sRNA Regulates RhlR-Dependent Quorum Sensing in Pseudomonas aeruginosa. mBio 10. 10.1128/mBio.02253-19.

57. Loh, E., Dussurget, O., Gripenland, J., Vaitkevicius, K., Tiensuu, T., Mandin, P., Repoila, F., Buchrieser, C., Cossart, P., and Johansson, J. (2009). A trans-Acting Riboswitch Controls Expression of the Virulence Regulator PrfA in Listeria monocytogenes. Cell 139, 770–779. 10.1016/j.cell.2009.08.046.

58. Ogura, M., Matsutani, M., Asai, K., and Suzuki, M. (2023). Glucose controls manganese homeostasis through transcription factors regulating known and newly identified manganese transporter genes in Bacillus subtilis. J Biol Chem 299, 105069. 10.1016/j.jbc.2023.105069.

59. Percy, M.G., and Gründling, A. (2014). Lipoteichoic acid synthesis and function in gram-positive bacteria. Annual review of microbiology 68, 81–100. 10.1146/annurev-micro-091213-112949.

60. Monk, I.R., Tree, J.J., Howden, B.P., Stinear, T.P., and Foster, T.J. (2015). Complete Bypass of Restriction Systems for Major Staphylococcus aureus Lineages. mBio 6, e00308–00315. 10.1128/mBio.00308-15.

61. Simons, R.W., Houman, F., and Kleckner, N. (1987). Improved single and multicopy lac-based cloning vectors for protein and operon fusions. Gene 53, 85–96.

62. Kehl-Fie, T.E., Zhang, Y., Moore, J.L., Farrand, A.J., Hood, M.I., Rathi, S., Chazin, W.J., Caprioli, R.M., and Skaar, E.P. (2013). MntABC and MntH contribute to systemic Staphylococcus aureus infection by competing with calprotectin for nutrient manganese. Infection and immunity 81, 3395–3405. 10.1128/iai.00420-13.

63. Arnaud, M., Chastanet, A., and Debarbouille, M. (2004). New vector for efficient allelic replacement in naturally nontransformable, low-GC-content, gram-positive bacteria. Appl Environ Microbiol 70, 6887–6891. 10.1128/aem.70.11.6887-6891.2004.

64. Wright, P.R., Georg, J., Mann, M., Sorescu, D.A., Richter, A.S., Lott, S., Kleinkauf, R., Hess, W.R., and Backofen, R. (2014). CopraRNA and IntaRNA: predicting small RNA targets, networks and interaction domains. Nucleic Acids Research 42, W119–123. 10.1093/nar/gku359.

65. Varadi, M., Anyango, S., Deshpande, M., Nair, S., Natassia, C., Yordanova, G., Yuan, D., Stroe, O., Wood, G., Laydon, A., et al. (2022). AlphaFold Protein Structure Database: massively expanding the structural coverage of protein-sequence space with high-accuracy models. Nucleic Acids Res 50, D439–d444. 10.1093/nar/gkab1061.

66. Meng, E.C., Goddard, T.D., Pettersen, E.F., Couch, G.S., Pearson, Z.J., Morris, J.H., and Ferrin, T.E. (2023). UCSF ChimeraX: Tools for structure building and analysis. Protein science : a publication of the Protein Society 32, e4792. 10.1002/pro.4792.

67. Koch, G., Yepes, A., Förstner, K.U., Wermser, C., Stengel, S.T., Modamio, J., Ohlsen, K., Foster, K.R., and Lopez, D. (2014). Evolution of resistance to a last-resort antibiotic in Staphylococcus aureus via bacterial competition. Cell 158, 1060–1071. 10.1016/j.cell.2014.06.046.

68. Schneider, C.A., Rasband, W.S., and Eliceiri, K.W. (2012). NIH Image to ImageJ: 25 years of image analysis. Nature methods 9, 671–675. 10.1038/nmeth.2089.

69. Lalaouna, D., Carrier, M.C., Semsey, S., Brouard, J.S., Wang, J., Wade, J.T., and Masse, E. (2015). A 3’ external transcribed spacer in a tRNA transcript acts as a sponge for small RNAs to prevent transcriptional noise. Molecular cell 58, 393–405. 10.1016/j.molcel.2015.03.013.

70. Salvail, H., Balaji, A., Roth, A., and Breaker, R.R. (2023). A spermidine riboswitch class in bacteria exploits a close variant of an aptamer for the enzyme cofactor S-adenosylmethionine. Cell Rep 42, 113571. 10.1016/j.celrep.2023.113571.

71. Boisset, S., Geissmann, T., Huntzinger, E., Fechter, P., Bendridi, N., Possedko, M., Chevalier, C., Helfer, A.C., Benito, Y., Jacquier, A., et al. (2007). Staphylococcus aureus RNAIII coordinately represses the synthesis of virulence factors and the transcription regulator Rot by an antisense mechanism. Genes Dev 21, 1353–1366. 10.1101/gad.423507.

72. Krute, C.N., Seawell, N.A., and Bose, J.L. (2021). Measuring Staphylococcal Promoter Activities Using a Codon-Optimized β-Galactosidase Reporter. Methods Mol Biol 2341, 37–44. 10.1007/978-1-0716-1550-8_6.

73. Battesti, A., Majdalani, N., and Gottesman, S. (2015). Stress sigma factor RpoS degradation and translation are sensitive to the state of central metabolism. Proc Natl Acad Sci U S A 112, 5159–5164. 10.1073/pnas.1504639112.

74. Majdalani, N., Cunning, C., Sledjeski, D., Elliott, T., and Gottesman, S. (1998). DsrA RNA regulates translation of RpoS message by an anti-antisense mechanism, independent of its action as an antisilencer of transcription. Proc Natl Acad Sci U S A 95, 12462–12467.

75. Kalvari, I., Nawrocki, E.P., Ontiveros-Palacios, N., Argasinska, J., Lamkiewicz, K., Marz, M., Griffiths-Jones, S., Toffano-Nioche, C., Gautheret, D., Weinberg, Z., et al. (2021). Rfam 14: expanded coverage of metagenomic, viral and microRNA families. Nucleic Acids Res 49, D192–d200. 10.1093/nar/gkaa1047.

76. Lott, S.C., Schäfer, R.A., Mann, M., Backofen, R., Hess, W.R., Voß, B., and Georg, J. (2018). GLASSgo - Automated and Reliable Detection of sRNA Homologs From a Single Input Sequence. Frontiers in genetics 9, 124. 10.3389/fgene.2018.00124.

77. Nawrocki, E.P., and Eddy, S.R. (2013). Infernal 1.1: 100-fold faster RNA homology searches. Bioinformatics 29, 2933–2935. 10.1093/bioinformatics/btt509.

78. Steinegger, M., and Söding, J. (2017). MMseqs2 enables sensitive protein sequence searching for the analysis of massive data sets. Nature biotechnology 35, 1026–1028. 10.1038/nbt.3988.

79. Sievers, F., Wilm, A., Dineen, D., Gibson, T.J., Karplus, K., Li, W., Lopez, R., McWilliam, H., Remmert, M., Söding, J., et al. (2011). Fast, scalable generation of high-quality protein multiple sequence alignments using Clustal Omega. Mol Syst Biol 7, 539. 10.1038/msb.2011.75.

80. Letunic, I., and Bork, P. (2024). Interactive Tree of Life (iTOL) v6: recent updates to the phylogenetic tree display and annotation tool. Nucleic Acids Res 52, W78–w82. 10.1093/nar/gkae268.

81. Kehl-Fie, T.E., Chitayat, S., Hood, M.I., Damo, S., Restrepo, N., Garcia, C., Munro, K.A., Chazin, W.J., and Skaar, E.P. (2011). Nutrient metal sequestration by calprotectin inhibits bacterial superoxide defense, enhancing neutrophil killing of Staphylococcus aureus. Cell host & microbe 10, 158–164. 10.1016/j.chom.2011.07.004.

82. Injarabian, L., Skerniskyte, J., Giai Gianetto, Q., Witko-Sarsat, V., and Marteyn, B.S. (2021). Reducing neutrophil exposure to oxygen allows their basal state maintenance. Immunology and cell biology 99, 782–789. 10.1111/imcb.12458.

