## Supplemental data for "An unprecedented small RNA-riboswitch interaction controls expression of a bifunctional pump that is essential for *Staphylococcus aureus* infection"

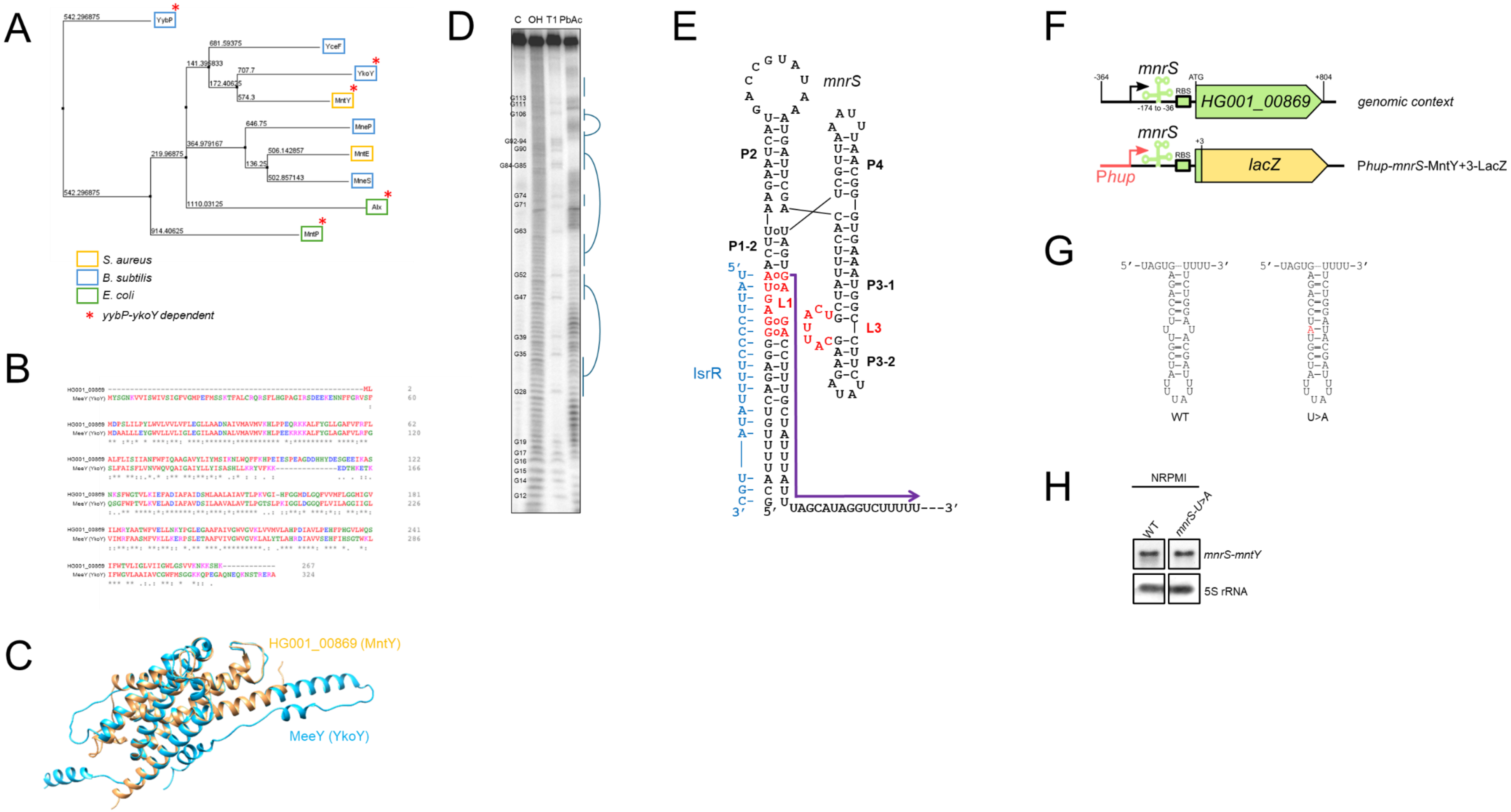
A
YybP
YceF
YkoY
MntY
MneP
MntE
MneS
Alx
MntP
S. aureus
B. subtilis
E. coli
* yybP-ykoY dependent
B
C
HG001_00869 (MntY)
MeeY (YkoY)
D
C OH T1 PbAc
E
mnrS
P2
P4
P1-2
P3-1
P3-2
L1
L3
IsrR
UAGCAUAGGUCUUUUU---3′
F
mnrS
HG001_00869
genomic context
lacZ
Phup
Phup-mnrS-MntY+3-LacZ
G
5′-UAGUG-UUUU-3′
WT
U>A
H
NRPMI
WT
mnrS-U>A
mnrS-mntY
5S rRNA

Figure S1

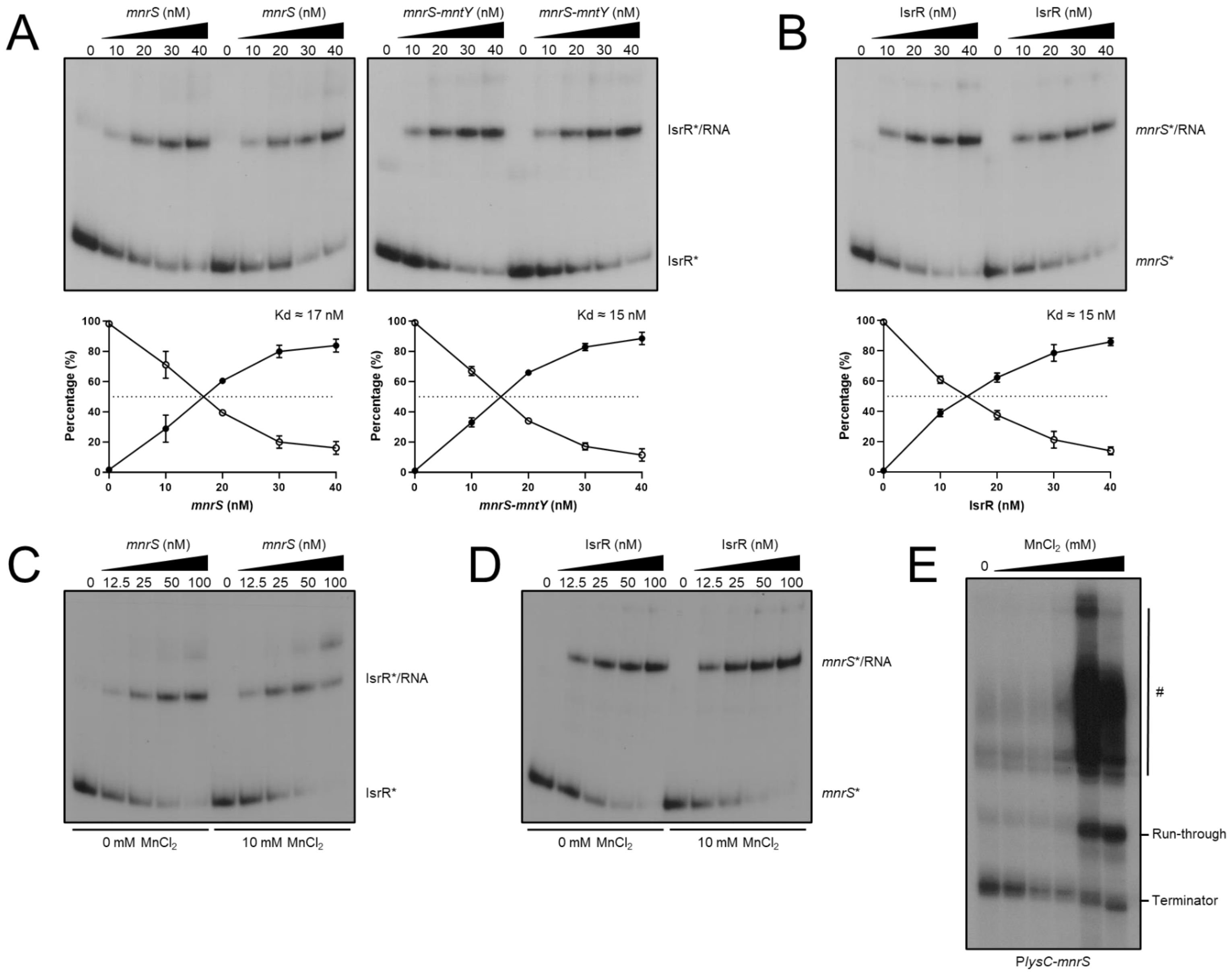
A
mnrS (nM)
0 10 20 30 40 0 10 20 30 40
mnrS-mntY (nM)
IsrR*/RNA
IsrR*
Kd ≈ 17 nM
Kd ≈ 15 nM
Percentage (%)
mnrS (nM)
mnrS-mntY (nM)
B
IsrR (nM)
mnrS*/RNA
mnrS*
IsrR (nM)
C
0 12.5 25 50 100 0 12.5 25 50 100
0 mM MnCl2
10 mM MnCl2
D
E
MnCl2 (mM)
0
#
Run-through
Terminator
PlysC-mnrS

Figure S2

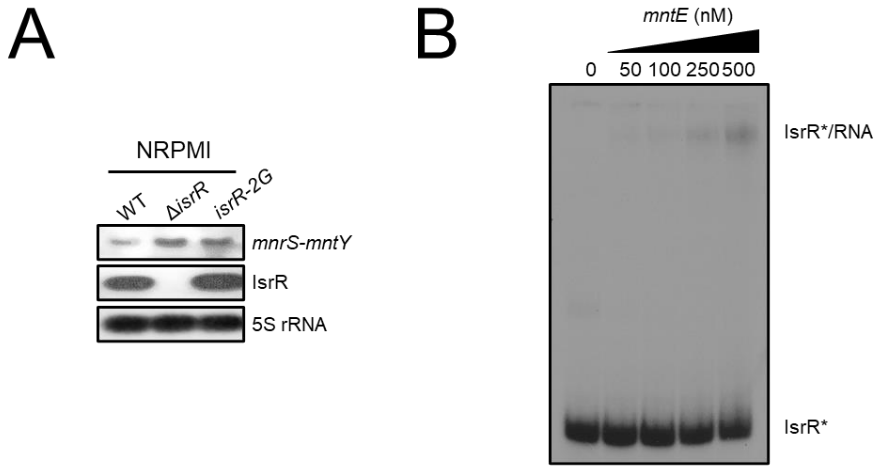
A
NRPMI
WT
ΔisrR
isrR-2G
mnrS-mntY
IsrR
5S rRNA
B
mntE (nM)
0
50
100
250
500
IsrR*/RNA
IsrR*

Figure S3

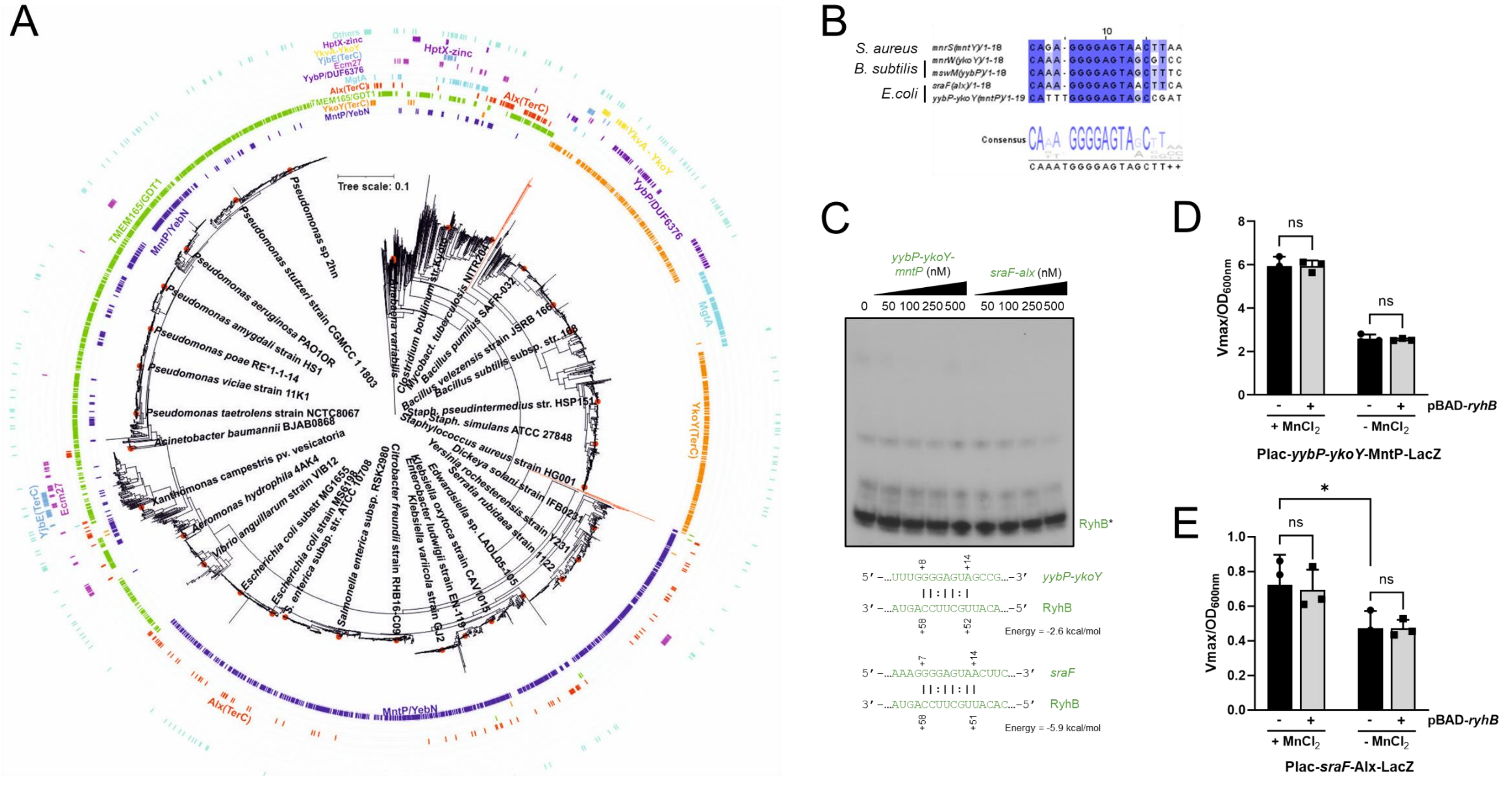

Figure S4

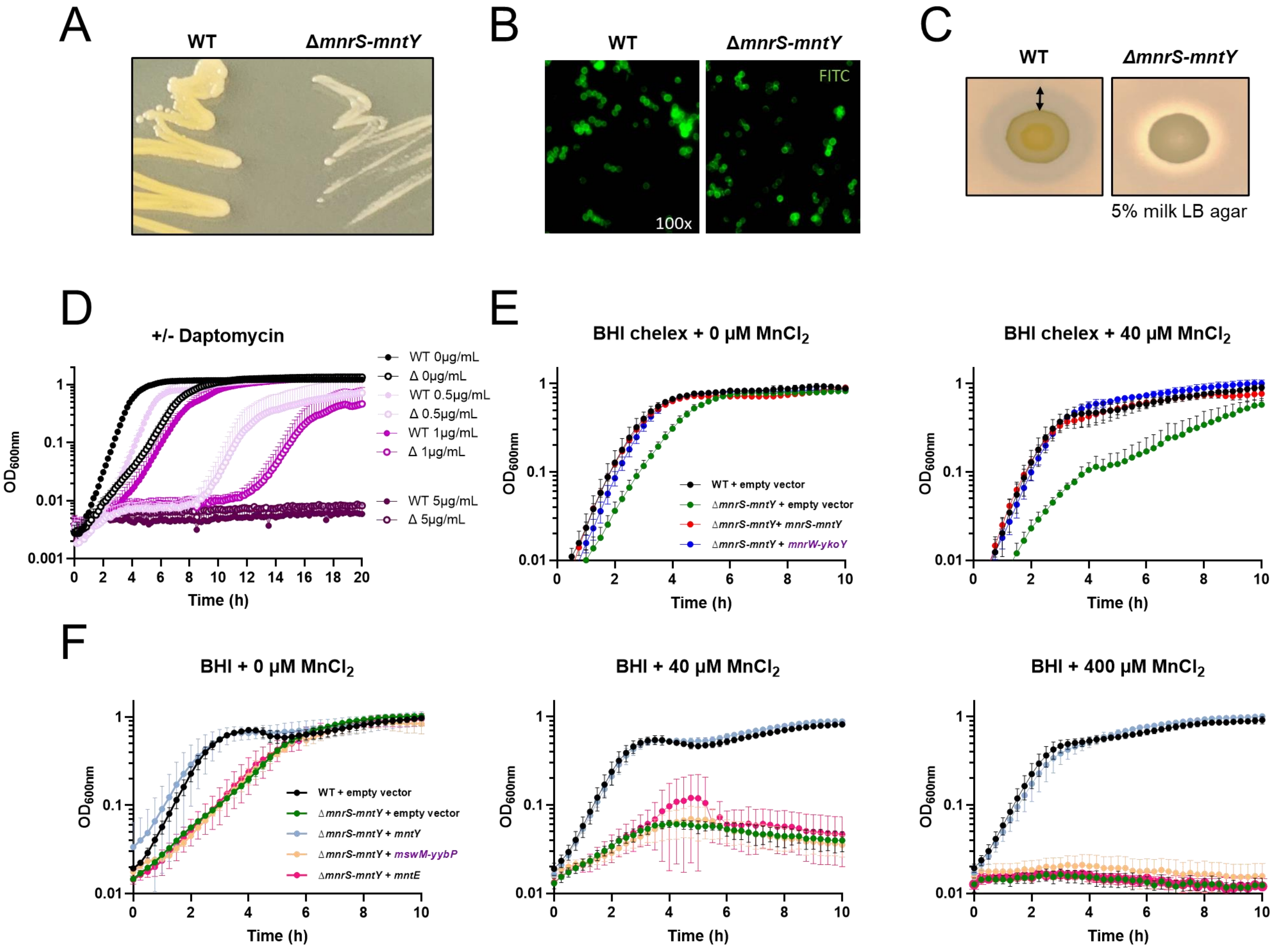

A
WT
ΔmnrS-mntY
B
WT
ΔmnrS-mntY
FITC
100x
C
WT
ΔmnrS-mntY
5% milk LB agar
D
+/- Daptomycin
WT 0μg/mL
Δ 0μg/mL
WT 0.5μg/mL
Δ 0.5μg/mL
WT 1μg/mL
Δ 1μg/mL
WT 5μg/mL
Δ 5μg/mL
OD600nm
Time (h)
E
BHI chelex + 0 μM MnCl2
BHI chelex + 40 μM MnCl2
WT + empty vector
ΔmnrS-mntY + empty vector
ΔmnrS-mntY+ mnrS-mntY
ΔmnrS-mntY + mnrW-ykoY
OD600nm
Time (h)
F
BHI + 0 μM MnCl2
BHI + 40 μM MnCl2
BHI + 400 μM MnCl2
WT + empty vector
ΔmnrS-mntY + empty vector
ΔmnrS-mntY + mntY
ΔmnrS-mntY + mswM-yybP
ΔmnrS-mntY + mntE
OD600nm
Time (h)


Figure S5

### SUPPLEMENTARY FIGURE LEGENDS

**Figure S1. *HG001_00869* codes for a putative Mn efflux pump and is under the control of the Mn-responsive *mnrS* riboswitch.** A. Multiple sequence alignment of proteins involved in Mn tolerance from *E. coli* (MntP and Alx, green boxes), *B. subtilis* (MneP, MneS, YkoY and YybP, blue boxes) and *S. aureus* (MntE and MntY, yellow boxes) was performed using Muscle (Output format: ClustalW; Edgar, 2004). Phylogenetic tree was reconstructed by the distance-based method "neighbor-joining" using Jalview (BLOSUM62; Procter et al., 2021). The presence of a *yybP-ykoY* motif upstream of the respective genes is indicated by a red star (*). B. Sequence alignment of HG001_00869 (*S. aureus* HG001; renamed MntY) and YkoY (*B. subtilis* 168; also named MeeY) proteins using ClustalW and default parameters. C. Superposition of AlphaFold models (Jumper et al., 2021) of HG001_00869 (*S. aureus* HG001 in yellow) and YkoY (*B. subtilis* in light blue) proteins using UCSF Chimera (Pettersen et al., 2014). D. Lead (PbAc) probing of 5'end-radiolabeled *mnrS* riboswitch (full-length) incubated in the absence of $MnCl_2$. C, non-reacted control; OH, alkaline ladder; T1, RNase T1 ladder. The numbers to the left show guanine (G) positions with respect to the transcription start (+1) of *mnrS*. Weakly cleaved regions are indicated with blue bars. Results are representative of at least two independent experiments. E. Secondary structure reflecting the 3D structure of the *mnrS* riboswitch. The conserved L1 and L3 loops are shown in red. Non-canonical base pairs are indicated by open circles. The terminator sequence is indicated by a purple arrow. The interaction between IsrR and the *mnrS* riboswitch (Figure 2B) is represented in blue. F. Description of P*hup-mnrS*-MntY+3 translational *lacZ* fusion used in this study. RBS, ribosome binding site; ATG, start codon. G. Secondary structure of the terminator of *mnrS* riboswitch. The mutation introduced to repair the terminator (U>A) in *S. aureus* is indicated in red. H. Northern blot analysis of *mnrS-mntY* RNA levels in WT and *mnrS*-U>A backgrounds in NRPMI medium only supplemented with 1 mM $MgCl_2$ and 100 µM $CaCl_2$. Cells were harvested at $OD_{600nm}$=1. 5S rRNA was used as loading control. Samples were loaded on the same gel. Results are representative of two independent experiments. Related to Figure 1.

**Figure S2. IsrR efficiently binds to *mnrS* riboswitch in vitro.** A. Gel retardation assays using IsrR sRNA (full-length), *mnrS* riboswitch (full-length) and *mnrS-mntY* mRNA (from -183 to +115). 5' end-radiolabeled IsrR (*) was incubated with increasing concentrations of *mnrS* or *mnrS-mntY* (0, 10, 20, 30 and 40 nM). The densitometric analyses were performed using ImageJ software (N=2). The percentage of free radiolabeled IsrR and IsrR-RNA complexes is indicated by white and black circles, respectively. In tested conditions, the Kd dissociation constant corresponds to the concentration of the cold RNA showing 50% of binding (dashed line). B. Gel retardation assays performed with 5' end-radiolabeled *mnrS* (*) and increasing concentrations of cold IsrR. C. Gel retardation assays using IsrR sRNA (full-length) and *mnrS* riboswitch (full-length). 5' end-radiolabeled IsrR (*) was incubated with increasing concentrations of *mnrS* (0, 12.5, 25, 50 and 100 nM) in the presence or absence of 10 mM $MnCl_2$. D. Gel retardation assays performed with 5' end-radiolabeled *mnrS* (*) and increasing concentrations of cold IsrR. E. Termination efficiency assays using P*lysC-mnrS* construct in the presence of increasing concentrations of $MnCl_2$ (0, 0.001, 0.01, 0.1, 1 and 10 mM). Results are representative of at least two independent experiments. # indicates artefact bands. Related to Figure 2.

**Figure S3. IsrR sRNA directly impairs the synthesis of full-length *mnrS-mntY*.** A. Northern blot analysis of *mnrS-mntY* RNA levels in WT, Δ*isrR* and *isrR-2G* mutant backgrounds; Cells were grown in NRPMI medium only supplemented with 1 mM $MgCl_2$ and 100 µM $CaCl_2$. 5S rRNA was used as loading control. Results are representative of two independent experiments. B. Gel retardation assays using IsrR sRNA (full-length) and *mntE mRNA* (full-length). The 5' end-radiolabeled IsrR (*) was incubated with increasing concentrations of *mntE* (0, 50, 100, 250 and 500 nM). Related to Figure 3.

**Figure S4. RyhB sRNA does not regulate the synthesis of MntP and Alx in *E. coli*, even if the pairing sequence on *yybP-ykoY* riboswitch is highly conserved.** A. 16S rDNA-based NJ showing the distribution of the detected riboswitch families. B. Multiple sequence alignment performed using ClustalW (Larkin et al., 2007) with default parameters and visualized using Jalview software (Procter et al., 2021). The 5'end sequence of *yybP-ykoY* riboswitches from *S. aureus* HG001 (*mnrS*(*mntY*)), *B. subtilis* 168 (*mswM*(*yybP*) and *mnrW*(*ykoY*)), and *E. coli* MG1655 (*yybP-ykoY*(*mntP*) and *sraF*(*alx*)) were extracted from RefSeq database (O'Leary et al., 2016). C. Gel retardation assays using RyhB (full-length), *yybP-ykoY-mntP* (from -225 to +50) and *sraF-alx* mRNA (from -206 to +170) from *E. coli* MG1655. The 5' end-radiolabeled RyhB sRNA (*) was incubated with increasing concentrations of cold mRNA (0, 50, 100, 250 and 500 nM). The putative pairing site predicted in silico using IntaRNA algorithm is indicated below (Wright et al., 2014). Results are representative of at least two independent experiments. D. β-galactosidase assays using *yybP-ykoY*-MntP+51-LacZ translational (in-frame) fusion under the control of the inductible pLac promoter. Δ*ryhB* mutant strains carry either the empty vector pNM12 (black bars (-)) or the pBAD-*ryhB* (gray bars (+)) were grown in LB medium in the presence of 1 mM IPTG. The expression of *ryhB* was induced by the addition of 0.1% arabinose when cells reached an $OD_{600nm}$ of 0.1. When indicated, 40 µM $MnCl_2$ were added. Samples were taken at an $OD_{600nm}$ of 0.8. Results are representative of at least three independent experiments ± SD. A two-way ANOVA analysis followed by Sidak's multiple comparison test was performed using Prism software (*p-value<0.05; ns, non-significant). E. β-galactosidase assays using *sraF*-Alx+15-LacZ translational (in-frame) fusion under the control of the inductible Plac promoter. See (D) for more details. Related to Figure 4.

**Figure S5. The deletion of *mnrS-mntY* induces strong phenotypic defects.** A. WT and Δ*mnrS-mntY* mutant strains streaked onto BHI agar plates and incubated at 37°C for 48h. B. Microscope observation of WT and Δ*mnrS-mntY* FITC-labelled cells. C. WT and Δ*mnrS-mntY* mutant strains streaked onto 5% skim milk LB agar plates. D. Growth monitoring ($OD_{600nm}$) of WT (closed circle) and Δ*mnrS-mntY* (Δ; open circle) strains in BHI medium supplemented with 0, 0.5, 1 or 5 µg/mL of daptomycin and 100 µg/mL $CaCl_2$ for 20h at 37°C. Data correspond to the mean of three independent experiments ± standard deviation (SD). E. Growth monitoring ($OD_{600nm}$) in chelex-treated BHI medium supplemented with 1 mM $MgCl_2$, 100 µM $CaCl_2$ ± 40 µM $MnCl_2$. WT and Δ*mnrS-mntY* strains contain the empty vector or a pEW derivate plasmid which enables the constitutive expression of *mnrS-mntY* or *mnrW-ykoY*. Gene sequences from *B. subtilis* are shown in purple. F. Growth monitoring ($OD_{600nm}$) in BHI medium supplemented with 0, 40 or 400µM $MnCl_2$ and 10 µg/mL erythromycin for 10h at 37°C. WT and Δ*mnrS-mntY* strains contain the empty vector or a pEW derivative plasmid, which enables the constitutive expression of *mntY*, *mntE* or *mswM-yybP*. Related to Figure 5.

### SUPPLEMENTARY TABLE LEGENDS

**Table S1. List of IsrR targets predicted by CopraRNA algorithm in this study.** Staphylococcal strains used for targets prediction are indicated in the section "Material and Methods". Each isrR gene sequence was extracted from RefSeq database (O'Leary et al., 2016). The cofactors associated to respective proteins were extracted from UniProt database (The UniProt Consortium, 2020). Previously validated mRNA targets are highlighted in gray.

**Table S2. Co-appearance and interaction potential of Fe-responsive sRNAs and Mn-sensing *yybP-ykoY* riboswitches in bacteria.**

**Table S3. Strains, plasmids and oligonucleotides used in this study.** A. List of strains, cell lines, mice and plasmids used in this study. B. List of oligonucleotides and gBlocks used in this study.

### SUPPLEMENTARY REFERENCES


1 Monk, I. R., Tree, J. J., Howden, B. P., Stinear, T. P. & Foster, T. J. Complete Bypass of Restriction Systems for Major Staphylococcus aureus Lineages. *mBio* **6**, e00308-00315, doi:10.1128/mBio.00308-15 (2015).

2 Huntzinger, E. *et al.* Staphylococcus aureus RNAIII and the endoribonuclease III coordinately regulate spa gene expression. *Embo j* **24**, 824-835, doi:10.1038/sj.emboj.7600572 (2005).

3 Arnaud, M., Chastanet, A. & Debarbouille, M. New vector for efficient allelic replacement in naturally nontransformable, low-GC-content, gram-positive bacteria. *Appl Environ Microbiol* **70**, 6887-6891, doi:10.1128/aem.70.11.6887-6891.2004 (2004).

4 Herbert, S. *et al.* Repair of global regulators in Staphylococcus aureus 8325 and comparative analysis with other clinical isolates. *Infection and immunity* **78**, 2877-2889, doi:10.1128/iai.00088-10 (2010).

5 Helle, L. *et al.* Vectors for improved Tet repressor-dependent gradual gene induction or silencing in Staphylococcus aureus. *Microbiology (Reading, England)* **157**, 3314-3323, doi:10.1099/mic.0.052548-0 (2011).

6 Masse, E. & Gottesman, S. A small RNA regulates the expression of genes involved in iron metabolism in Escherichia coli. *Proc Natl Acad Sci U S A* **99**, 4620-4625, doi:10.1073/pnas.032066599 (2002).

7 Majdalani, N., Cunning, C., Sledjeski, D., Elliott, T. & Gottesman, S. DsrA RNA regulates translation of RpoS message by an anti-antisense mechanism, independent of its action as an antisilencer of transcription. *Proc Natl Acad Sci U S A* **95**, 12462-12467 (1998).

8 Krute, C. N., Seawell, N. A. & Bose, J. L. Measuring Staphylococcal Promoter Activities Using a Codon-Optimized β-Galactosidase Reporter. *Methods Mol Biol* **2341**, 37-44, doi:10.1007/978-1-0716-1550-8_6 (2021).

9 Consortium, T. U. UniProt: the universal protein knowledgebase in 2021. *Nucleic Acids Research* **49**, D480-D489, doi:10.1093/nar/gkaa1100 (2020).
10 Larkin, M. A. *et al.* Clustal W and Clustal X version 2.0. *Bioinformatics* **23**, 2947-2948, doi:10.1093/bioinformatics/btm404 (2007).
11 Procter, J. B. *et al.* Alignment of Biological Sequences with Jalview. *Methods Mol Biol* **2231**, 203-224, doi:10.1007/978-1-0716-1036-7_13 (2021).
12 O'Leary, N. A. *et al.* Reference sequence (RefSeq) database at NCBI: current status, taxonomic expansion, and functional annotation. *Nucleic Acids Res* **44**, D733-745, doi:10.1093/nar/gkv1189 (2016).
13 Pettersen, E. F. *et al.* UCSF Chimera--a visualization system for exploratory research and analysis. *Journal of computational chemistry* **25**, 1605-1612, doi:10.1002/jcc.20084 (2004).
14 Menendez-Gil, P. *et al.* Differential evolution in 3'UTRs leads to specific gene expression in Staphylococcus. *Nucleic Acids Res* **48**, 2544-2563, doi:10.1093/nar/gkaa047 (2020).
15 Nair, D. *et al.* Whole-genome sequencing of Staphylococcus aureus strain RN4220, a key laboratory strain used in virulence research, identifies mutations that affect not only virulence factors but also the fitness of the strain. *J Bacteriol* **193**, 2332-2335, doi:10.1128/jb.00027-11 (2011).
16 Bae, T. & Schneewind, O. Allelic replacement in Staphylococcus aureus with inducible counter-selection. *Plasmid* **55**, 58-63, doi:10.1016/j.plasmid.2005.05.005 (2006).
17 Desnoyers, G., Morissette, A., Prevost, K. & Masse, E. Small RNA-induced differential degradation of the polycistronic mRNA iscRSUA. *EMBO J* **28**, 1551-1561, doi:10.1038/emboj.2009.116 (2009).
18 Simons, R. W., Houman, F. & Kleckner, N. Improved single and multicopy lac-based cloning vectors for protein and operon fusions. *Gene* **53**, 85-96 (1987).
19 Jumper, J. *et al.* Highly accurate protein structure prediction with AlphaFold. *Nature* **596**, 583-589, doi:10.1038/s41586-021-03819-2 (2021).
20 Edgar, R. C. MUSCLE: multiple sequence alignment with high accuracy and high throughput. *Nucleic Acids Res* **32**, 1792-1797, doi:10.1093/nar/gkh340 (2004).
